# Rapid turnover of sex-determining architecture in invasive mussels

**DOI:** 10.64898/2026.03.18.712737

**Authors:** Alexandra A.-T. Weber, Kavitha Uthanumallian, Kevin M. Kocot, Marco Giulio, Silvia G. Signorini, Marie-Claude Senut, Zeyuan Chen, Julia D. Sigwart, Yale J. Passamaneck

## Abstract

Sex determination is a fundamental developmental switch, yet the genetic architectures that underlie transitions between sex-determining systems remain poorly understood, especially in non-model animals with homomorphic sex chromosomes. Here, we combine chromosome-scale genomes, whole-genome resequencing of 80 sexed individuals, sex-specific k-mer analyses, and developmental transcriptomics to resolve and compare sex determination in zebra and quagga mussels, two closely related and globally invasive freshwater bivalves. We find that these species have evolved sharply contrasting sex-determining architectures since their divergence. Zebra mussels show signatures of a polygenic ZZ/ZW system, with sex-linked regions on multiple chromosomes, including signal at the conserved ovarian regulator *FoxL2*. In contrast, quagga mussels possess a localized ∼800 kb XX/XY sex-determining region containing *FoxL2-Y*, a duplicated and structurally modified *FoxL2* paralog and candidate male-determining locus. In quagga mussels, SNP-based differentiation, model-based inference, sex-biased read coverage, and male-specific k-mers independently converge on this Y-linked region, while diagnostic k-mer screening of staged embryonic transcriptomes reveals rare, early developmental expression of *FoxL2-Y*, consistent with a transient regulatory role. The quagga sex-determining region is enriched for C-type lectins and other genes implicated in reproductive interactions, suggesting that haploid selection may have contributed to the stabilization of this region. Together, these findings show that closely related species can rapidly evolve different genetic solutions to sex determination and identify gene duplication, genome dynamism, and reproductive selection as potential contributors to sex-determining system turnover. Because zebra and quagga mussels are major freshwater invaders, resolving their divergent reproductive architectures also provides essential genomic context for assessing the future feasibility of sex-ratio-based biocontrol strategies.

## Introduction

Biological invasions are among the most consequential global environmental changes of the Anthropocene, reshaping ecosystems, altering food webs, and imposing substantial economic costs. Among freshwater invaders, the zebra mussel (*Dreissena polymorpha*; Pallas, 1771) and the quagga mussel (*Dreissena rostriformis bugensis*; Andrusov, 1897) stand out as two of the most ecologically and economically damaging species worldwide (*1*–*3*). Native to the Ponto–Caspian region, both species were introduced independently to North America and Western Europe during the last century, and their spread is still ongoing (*4*). Their extraordinary invasion success is driven by high fecundity, broadcast spawning, broad ecological tolerance, and the ability to reach extreme population densities, resulting in profound impacts on ecosystem functioning, native biodiversity, and water infrastructure, with substantial long-term management costs (*5*). Consequently, the zebra mussel has been listed among the world’s 100 worst invasive alien species (*6*). Despite decades of research, there are currently no efficient or scalable methods to control or eradicate quagga or zebra mussels once they have been established in natural ecosystems (*7*). This limitation has prompted growing interest in genetic biocontrol approaches, which aim to suppress, manage, or potentially eradicate invasive populations by manipulating their genetic or reproductive systems (*8*). In freshwater mussels, several conceptual and experimental efforts are underway to explore the feasibility of such approaches (*9*). One proposed strategy is to disrupt reproduction by biasing population sex ratios through the release of genetically modified individuals (*8*). In principle, recent advances in CRISPR–Cas9–based gene drive technologies could enable sex-ratio distortion in invasive mussels, potentially leading to population suppression or collapse (*7*, *9*). A critical prerequisite for any such strategy, however, is a detailed understanding of the underlying sex determination (SD) system—that is, the genetic and developmental mechanisms that specify male and female phenotypes. To date, the SD systems of zebra and quagga mussels have remained entirely uncharacterized.

Across eukaryotes, sex determination exhibits remarkable diversity, ranging from highly conserved systems to extreme evolutionary lability (*10*, *11*). In gonochoric species with separate sexes, genetic sex determination (GSD) mediated by sex chromosomes carrying a sex-determining gene is perhaps the most familiar mechanism, exemplified by the XX–XY system in mammals or the ZZ–ZW system in birds (*10*). However, sex can also be determined by the combined effects of multiple loci distributed across the genome, a configuration known as polygenic sex determination (PSD) (*12*–*14*). In other taxa, environmental cues such as temperature play a decisive role, as in many reptiles (*10*). Increasingly, evidence suggests that genetic and environmental influences can interact, giving rise to condition-dependent sex determination systems (*15*). These diverse mechanisms underscore that sex determination is not a static trait, but a developmental process that can evolve rapidly under appropriate ecological and genomic conditions. In bivalves, available evidence indicates that both genetic and environmental factors may contribute to sex determination (*16*, *17*). Yet, despite the group’s enormous diversity—over 18,000 described species—detailed SD studies exist for only a handful of taxa, largely focused on marine species of commercial interest such as oysters, scallops, and marine mussels (*17*). Cytogenetic data are available for approximately 150 bivalve species, all of which possess homomorphic chromosomes that are morphologically indistinguishable between males and females (*16*). This pattern suggests that bivalve SD may involve young or cryptic sex chromosomes, polygenic architectures, environmental effects, or combinations thereof. Importantly, homomorphic sex chromosomes are not necessarily evolutionarily young: recent work in scallops has revealed ancient homomorphic sex chromosomes involving conserved female-associated genes such as *FoxL2*, *ZNF226*, and *CYP3A24* (*18*). Across bivalves, genes of the *DMRT*, *Sox*, and *Fox* families frequently show sex-biased expression and have repeatedly been implicated in sex-determining pathways, in particular the bivalve-specific gene *DMRT-1L* and the gene *FoxL2*, for male and female pathways, respectively (*17*, *19*–*22*). A recent comparative genomic survey of 24 bivalve species revealed that *DMRT-1L* is absent from both zebra and quagga mussel genomes (*23*), consistent with earlier genome assemblies (*24*, *25*). Moreover, *DMRT-2* is absent from the zebra mussel genome, and the remaining *DMRT* paralogs (*DMRT-2*, *DMRT-3*, *DMRT-4/5*) show little evidence of accelerated evolution across bivalves (*23*), suggesting that they are unlikely to function as primary sex-determining genes in these species. Notably, however, within-species male–female genomic differentiation at these loci has received little attention, and population-level analyses of sex-determining gene families remain scarce in bivalves in general and absent for *Dreissena*.

Zebra and quagga mussels are gonochoric broadcast spawners with external fertilization, pelagic larval development, and benthic settlement, resembling the canonical reproductive cycle of marine molluscs (*3*). Zebra mussels possess 16 homomorphic chromosomes (*26*), a number mirrored by the 16 linkage groups in the available chromosome-scale reference genome (*25*). In contrast, no karyotype data are available for quagga mussels, and the published reference genome remains highly fragmented (>18,000 scaffolds) (*24*), severely limiting the identification of physically linked sex-determining loci. This limitation is not unique to *Dreissena*: across Mollusca, chromosome-scale reference genomes have historically been scarce, reflecting longstanding challenges associated with assembling large, repeat-rich, and highly heterozygous genomes in this hyper-diverse phylum (*27*). As a result, chromosome-level assemblies are essential to resolve sex-linked genomic regions, reconstruct coevolving gene blocks, and enable meaningful comparative analyses of sex-determining architectures. The evolutionary context of the genus *Dreissena* further motivates such comparisons. Originating in Lake Pannon—an extinct lake that existed from approximately 12 to 4 million years ago—the genus diversified rapidly during the late Miocene (*28*, *29*). Fossil and molecular evidence indicate that the zebra mussel (subgenus *Dreissena* s. str.) and the quagga mussel (subgenus *Pontodreissena*) diverged shortly after the formation of Lake Pannon, around 10–12 million years ago (*30*–*32*). While sex determination can remain highly conserved over deep evolutionary timescales, as in mammals and birds, it can also undergo rapid turnover, as documented in fishes and reptiles (*10*). Whether zebra and quagga mussels share a conserved SD system or have evolved distinct mechanisms since their divergence has remained an open question.

Population genomic approaches provide powerful tools to identify sex-linked loci in non-model organisms, particularly when sex chromosomes are homomorphic (*33*). By leveraging sex-specific differences in allele frequencies, heterozygosity, and sequencing coverage, whole-genome resequencing can reveal sex-linked regions even in the absence of visible chromosomal differentiation. Recent methodological advances, including tools such as RADsex (*34*), detsex (*35*), Findzx (*36*), and SDpop (*37*), have enabled the discovery of sex chromosomes and sex-linked loci across a wide range of taxa (*38*, *39*), opening new avenues for studying SD evolution in previously intractable systems. Here, we combine chromosome-scale genomics, population resequencing, and transcriptomic analyses to resolve the sex determination systems of two of the world’s most successful freshwater invaders. We generated a chromosome-scale reference genome and karyotype data for the quagga mussel, and used whole-genome resequencing of male and female zebra and quagga mussels to (i) identify sex-linked genomic regions in each species and (ii) compare their SD architectures in an explicit evolutionary framework. Finally, we integrate adult tissue-specific and early developmental RNA sequencing to characterize expression patterns of candidate loci across tissues and developmental stages in quagga mussels. By revealing fundamentally different SD architectures in two closely related invasive species, our study illuminates how dynamic genomes can facilitate rapid divergence of reproductive systems and provides critical genomic context for evaluating the feasibility of sex-based genetic biocontrol strategies in non-model invasive taxa.

## Results

### Many genomic rearrangements between the quagga and zebra mussel genomes

We first generated a chromosome-scale reference genome of the quagga mussel (Fig. 1A) to compare genomic rearrangements and the genetic architecture of sex-linked genes in both species, and to provide a high-quality genomic resource for quagga mussel research. Briefly, a total of 308 Gbp of PacBio HiFi data and 200 Gbp of Illumina paired-end (PE) data were generated from a single male specimen (∼317x coverage). GenomeScope analysis of the quality- and adapter-trimmed and filtered Illumina PE reads resulted in a predicted genome size of 1.08 Gbp and a heterozygosity of 2.45%. The initial genome assembly consisted of 8960 contigs totaling 2.84 Gbp with an N50 of 770 Mbp and an L50 of 871, significantly larger than the size predicted by GenomeScope. However, given the very high estimated heterozygosity, purging of duplicate haplotigs and polishing reduced the assembly to 2714 contigs totaling 1.61 Gbp with an N50 of 1.53 Mbp and an L50 of 311. Scaffolding of the haplotig-purged and polished assembly using Hi-C data yielded 16 chromosome-scale pseudomolecules and included 95.7% of the bases in the final pre-scaffolding assembly (Fig. 1B). Karyotype analysis confirmed that quagga mussels have 16 pairs of chromosomes (Fig. 1C).

**Figure 1:**
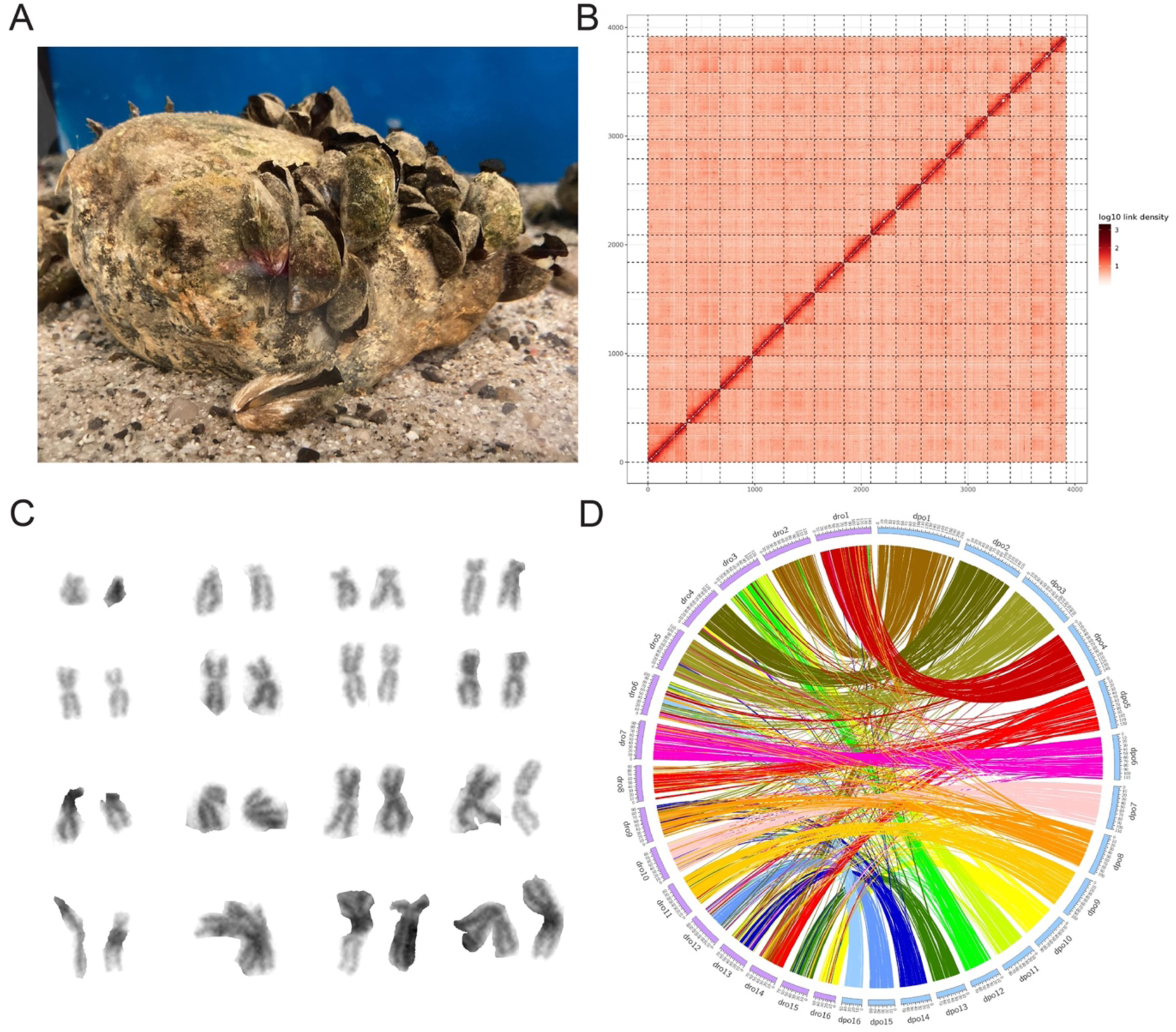
A new chromosome-scale genome assembly for the quagga mussel *Dreissena rostrifomis bugensis*. **A)** Illustrative picture of quagga mussels. **B)** Post-scaffolding heatmap of the present genome assembly. 95% of the scaffolds are placed on 16 linkage groups. **C)** Karyogram of the quagga mussel showing 16 pairs of chromosomes. **D)** Whole-genome alignment of the quagga mussel and the zebra mussel genomes show major rearrangements on several linkage groups. dro: *Dreissena rostriformis bugensis*. dpo: *Dreissena polymorpha*.

BUSCO analysis of the 16 Hi-C scaffolds detected 96.2% of the metazoa_odb10 single-copy orthologs as complete (92.6% single-copy, 2.1% duplicated, and 1.5% fragmented). RepeatMasker annotated 37.9% of the genome (611.9 Mb) as interspersed repetitive sequence. Unclassified repeats constitute the majority (27.7%) with classified transposable elements including LINEs (4.60%), DNA transposons (3.50%), LTR retrotransposons (1.49%), and SINEs (0.68%). Additional repetitive sequences include simple repeats (2.21%) with smaller contributions from low-complexity regions (0.09%) and satellites (0.01%). Genome annotation of the Hi-C scaffolds with BRAKER resulted in 83,983 predicted gene models with 97.8% of BUSCOs detected (90.3% single, 5.2% duplicated, 2.3% fragmented). Removal of gene models not supported by any transcriptome or protein evidence resulted in a final set of 58,582 gene models with 97.6% of BUSCOs detected (90.1% single, 5.2% duplicated, 2.3% fragmented).

We performed a synteny analysis between the newly assembled quagga mussel reference genome and the publicly available zebra mussel reference genome (*25*). Both species have similar genome sizes (∼1.6 Gbp) and the same chromosome number (16), which led us to anticipate a relatively conserved chromosomal structure with few major rearrangements. However, our analysis revealed notable genomic differences. While some chromosomes were highly conserved between the species, many others exhibited substantial rearrangements, underscoring the dynamic nature of both genomes (Fig. 1D). For example, linkage group (LG) 13 and LG14 in quagga mussels appear to have undergone extensive rearrangements, as they do not have clear counterparts in the zebra mussel chromosomes. In contrast, several chromosomes showed fewer rearrangements and appear to be largely conserved between the two species, such as LG1 of the quagga mussel and LG4 of the zebra mussel, or LG4 of the quagga mussel and LG2 of the zebra mussel. Finally, a large-scale fission/fusion event appears to have occurred between LG3 of quagga mussels and LG11 and LG12 of zebra mussels, although the directionality of this rearrangement—whether LG3 of quagga mussels resulted from the fusion of LG11 and LG12 of zebra mussels or vice versa—remains unclear due to the lack of an ancestral karyotype.

### Genomic variation and absence of differentiated sex chromosomes in quagga and zebra mussels

To uncover potential sex-linked loci in both species, we then generated whole-genome resequencing data of 20 males and 20 females from each species (Table 1, Table S1). The resequencing data was aligned to the respective reference genome of each species (both from a male specimen). Since our study presents the first whole-genome resequencing data for both invasive species, we first briefly describe patterns of genetic variation in these populations. Quagga mussels had a higher number of filtered variants than zebra mussels (Table 1), and genome-wide heterozygosity of the reference genome was likewise higher in quagga mussels. We then compared genomic differentiation between males and females of both species. We found that on a genome-wide level, there was no genetic differentiation between males and females of both species as highlighted by the male-female *F*_ST_ values close to zero (Table 1). These results highlight the absence of differentiated sex chromosomes in both quagga and zebra mussels.

**Table 1:** Diversity statistics and genome-wide male-female differentiation in zebra and quagga mussels.

| Statistics | Zebra mussel:<br>*Dreissena polymorpha* | Quagga mussel:<br>*Dreissena rostriformis bugensis* |
| --- | --- | --- |
| Number of filtered SNPs | 24,499,661 | 42,073,009 |
| Genome-wide heterozygosity<br>(from reference genome) | 2.13% | 2.45% |
| Genome-wide male-female $F_{ST}$<br>(W&C weighted $F_{ST}$) | 0.00054 | -0.00034 |
| mean depth in females | 8.4 | 9.6 |
| mean depth in males | 9.8 | 9.8 |

### Genomic signatures of polygenic ZZ-ZW sex determining system in zebra mussels

We then investigated local male–female differentiation using SNP-based *F_ST_* scans across the genome. In zebra mussels, several regions showed *F_ST_* ≈ 0.5 (Fig. 2A), consistent with the expected value for fully differentiated sex-linked SNPs under the Weir and Cockerham estimator (*40*, *41*). Interestingly, there was a region with very high male-female differentiation (*F*_ST_ ≈ 0.9) on LG13, as well as additional regions with *F*_ST_ >0.5 (Fig. 2A). To uncover which SD system is most likely to occur in zebra mussels, we highlighted ZW SNPs in pink (Fig. 2A) and XY SNPs in green (Fig. S1A) on the genome-wide *F*_ST_ plots. We found that zebra mussels likely possess a polygenic ZZ-ZW SD system, with at least three chromosomes (LG5, LG9, LG13) displaying signatures of a ZZ-ZW system with ZW SNPs colocalizing on *F*_ST_ peaks. We found very similar patterns of genomic differentiation when performing GWAS analyses, which also allowed us to remove some noise inherent to *F*_ST_ calculations (Fig. S2A-D). We then conducted a model-based approach – SDpop - to specifically test the likelihood of a XX-XY system, ZZ-ZW system and the NULL model (no sex chromosome) along non-overlapping 10 kb windows (*37*). We plotted the difference in Bayesian information criterion - BIC (a measure of model fit) between the XY and NULL model, and between the ZW and NULL model, where a large BIC difference (>2000) indicates a higher likelihood of the specific sex model (ZW or XY) (Fig. 2B). The SDpop results were congruent with *F*_ST_ and ZW SNPs for regions localizing on LG5 (Fig. 2D) and LG13 (Fig. 2H), while we did not find a SDpop signal on LG9 (Fig. 2F), contrary to a clear *F*_ST_ and ZW SNP peak (Fig. 2E). Despite the absence of SDpop signal on LG9, we decided to retain the region on LG9 as a candidate for further exploration. From these results, we identify eight candidate ZW-linked regions on three linkage groups in zebra mussels: three on LG5, one on LG9 and four on LG13 (Table 2).

**Figure 2:**
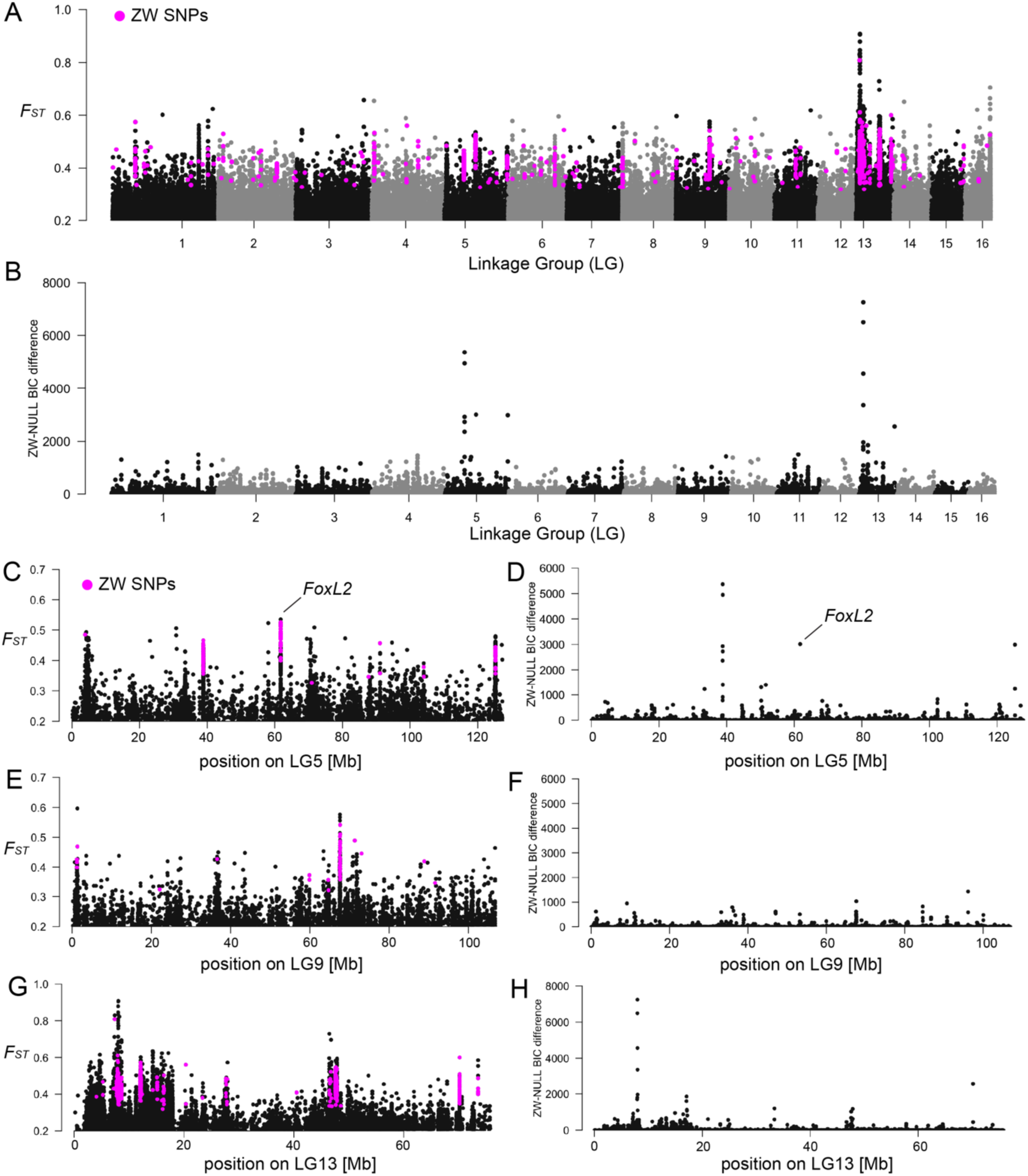
Genomic signatures of a polygenic ZZ-ZW sex determining system in the zebra mussel *Dreissena polymorpha*. **A)** SNP-based whole-genome Weir & Cockerham *F*_ST_ between 20 male and 20 female zebra mussels. ZW SNPs (homozygous in males and heterozygous in females) are highlighted in pink. Note that the Y axis starts at 0.2 for better readability in all *F*_ST_ plots. **B)** Bayesian Information Criterion (BIC) difference between ZW SDpop model and NULL model calculated over 10 kb sliding windows. An elevated BIC difference (>2000) corresponds to more support for the ZW model (ZZ-ZW sex determining system) compared with the NULL model (no genomic differentiation between males and females) for a particular 10 kb window. **C)** Male-female genome-wide *F*_ST_ for linkage group 5 (CM035919.1). The position of the female sex-determining gene *FoxL2* is highlighted. **D)** BIC differences between ZW and NULL SDpop model for linkage group 5 (CM035919.1). The position of the female sex-determing gene *FoxL2* is highlighted. **E)** Male-female genome-wide *F*_ST_ for linkage group 9 (CM035923.1). **F)** BIC differences between ZW and NULL SDpop model for linkage group 9 (CM035923.1). **G)** Male-female genome-wide *F*_ST_ for linkage group 13 (CM035927.1). **H)** BIC differences between ZW and NULL SDpop model for linkage group 13 (CM035927.1).

**Table 2:**
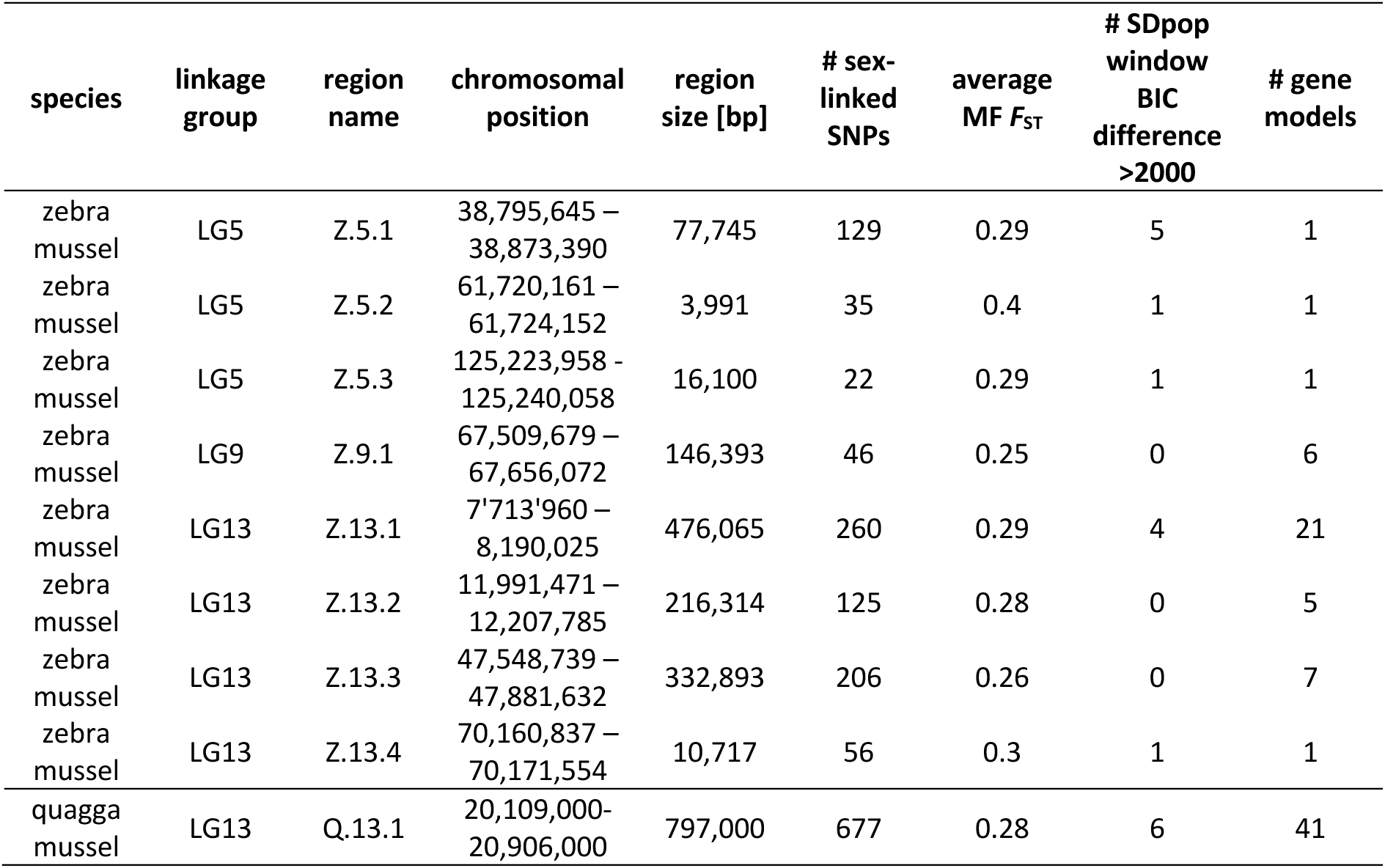
characteristics of candidate sex-linked regions in zebra (ZW) and quagga (XY) mussels.

| species | linkage group | region name | chromosomal position | region size [bp] | # sex-linked SNPs | average MF $F_{ST}$ | # SDpop window BIC difference >2000 | # gene models |
| --- | --- | --- | --- | --- | --- | --- | --- | --- |
| zebra mussel | LG5 | Z.5.1 | 38,795,645 – 38,873,390 | 77,745 | 129 | 0.29 | 5 | 1 |
| zebra mussel | LG5 | Z.5.2 | 61,720,161 – 61,724,152 | 3,991 | 35 | 0.4 | 1 | 1 |
| zebra mussel | LG5 | Z.5.3 | 125,223,958 – 125,240,058 | 16,100 | 22 | 0.29 | 1 | 1 |
| zebra mussel | LG9 | Z.9.1 | 67,509,679 – 67,656,072 | 146,393 | 46 | 0.25 | 0 | 6 |
| zebra mussel | LG13 | Z.13.1 | 7,713,960 – 8,190,025 | 476,065 | 260 | 0.29 | 4 | 21 |
| zebra mussel | LG13 | Z.13.2 | 11,991,471 – 12,207,785 | 216,314 | 125 | 0.28 | 0 | 5 |
| zebra mussel | LG13 | Z.13.3 | 47,548,739 – 47,881,632 | 332,893 | 206 | 0.26 | 0 | 7 |
| zebra mussel | LG13 | Z.13.4 | 70,160,837 – 70,171,554 | 10,717 | 56 | 0.3 | 1 | 1 |
| quagga mussel | LG13 | Q.13.1 | 20,109,000–20,906,000 | 797,000 | 677 | 0.28 | 6 | 41 |

### Candidate sex-determining genes in zebra mussels

There were three candidate SD regions on LG5 (Z.5.1-Z.5.3) with congruent sex-determining signatures (*F*_ST_ ≈ 0.5; ZW SNPs; SDpop signal) (Fig. 2C). These small regions encompassed a single gene model each (Table 2, Table S2), including one encoding the female sex-determining gene *FoxL2* (region Z.5.2). The gene model in region Z.5.1 is an uncharacterized protein with a putative metal ion binding function. BlastP results indicated that this gene encodes for a protein that possesses a tudor domain (Table S2). The gene model on region Z.5.3 (nucleolar complex protein 2 homolog) should be involved in negative regulation of transcription by RNA polymerase II.

The candidate SD region on LG9 is about 150 kb and contains six gene models (Table 2). Among these, one uncharacterized gene contains C2H2-type Zinc finger domains and shows homology to DNA-binding transcription factors with RNA polymerase II-specific activity (Table S2). Finally, LG13 was the chromosome that showed the highest signature of male-female genomic differentiation. We identified four candidate sex-linked regions based on the congruence of the different methods. The first region Z.13.1 is about 476 kb and contains 21 gene models (Table 2). It is the region that has the clearest SDpop signal across the full LG13 chromosome (Fig. 2H). Many gene models in this region are uncharacterized proteins, and five of them encode for long non-coding RNAs (Table S2). The second and the third regions do not have a strong SDpop signal (no window with BIC difference > 2000), however we believe it is of interest to discuss them given the congruence of male-female *F*_ST_ and the presence of a tight cluster of ZW SNPs (Fig. 2G). The second region Z.13.2 is about 216 kb and contains five gene models, including two long non-coding RNA (Table S2). The third region Z.13.3 is about 332 kb and contains seven gene models, including a RNA polymerase II transcription factor (Table S2). Finally, the fourth region Z.13.4 is about 10 kb and contains a single gene model encoding for a long non-coding RNA (Table S2).

### Localized genomic signature of an XX–XY sex-determining system in quagga mussels

We applied the same genome-wide approach used for zebra mussels to quagga mussels, performing male–female *F*_ST_ analyses across the genome (Fig. 3). In contrast to the diffuse patterns observed in zebra mussels, all sex-linked signals in quagga mussels converged on a single region on LG13, where elevated male–female *F*_ST_, clusters of sex-linked SNPs, and a strong SDpop signal colocalized (Fig. 3A–D). The absence of ZW-type SNP clustering and the lack of support for a ZW model in SDpop analyses (Fig. S3) indicate that quagga mussels most likely exhibit an XX–XY sex-determination system. This defines a single, relatively large (∼800 kb) candidate sex-determining (SD) region on LG13 (Q.13.1; Table 2), which was also identified using a GWAS analysis (Fig. S4).

**Figure 3:**
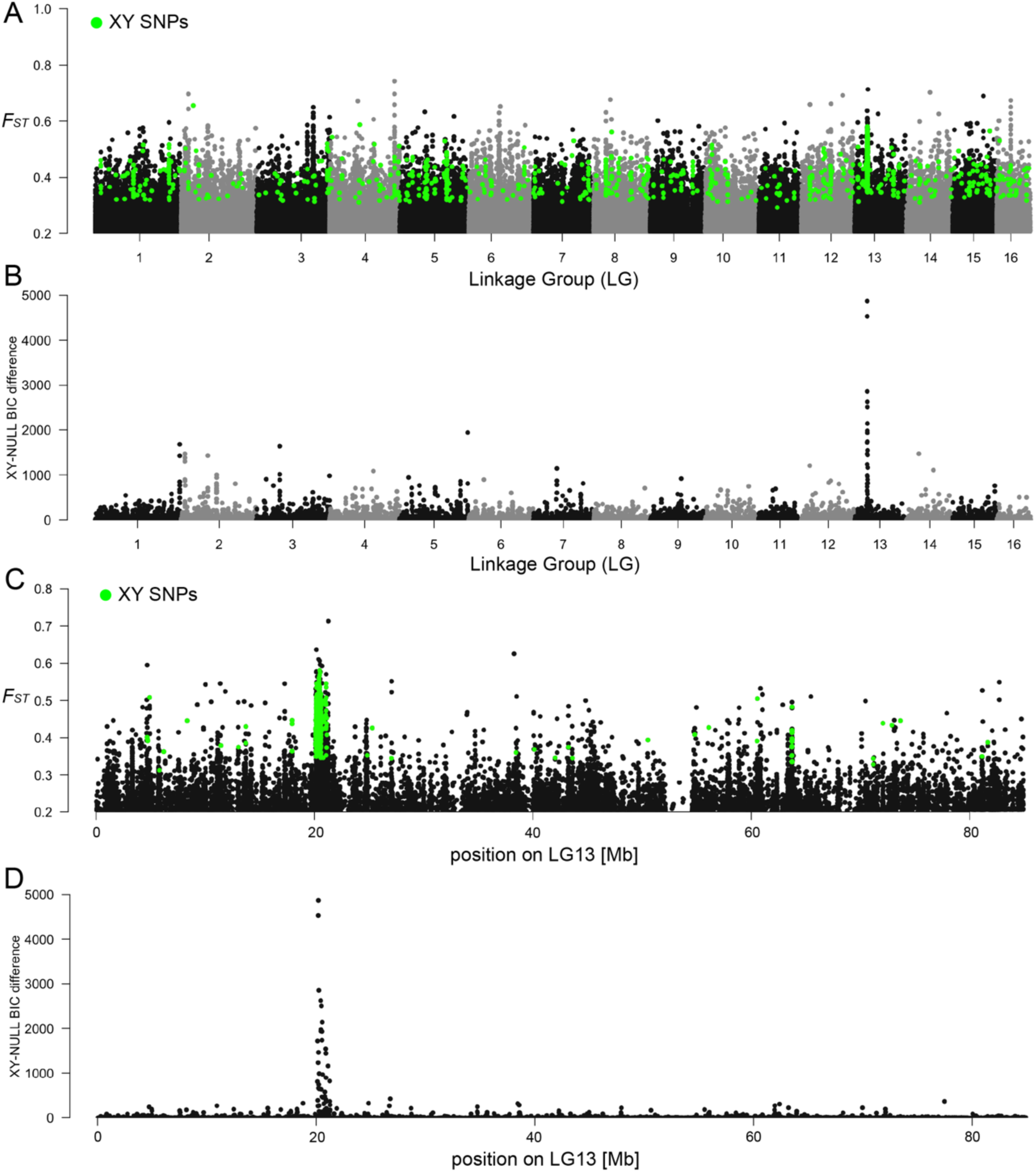
Genomic signatures of a localized XX-XY sex determining system in the quagga mussel *Dreissena rostriformis bugensis*. **A)** SNP-based whole-genome Weir & Cockerham *F*_ST_ between 19 male and 18 female quagga mussels. XY SNPs (heterozygous in males and homozygous in females) are highlighted in green. Note that the Y axis starts at 0.2 for better readability in all *F*_ST_ plots. **B)** Bayesian Information Criterion (BIC) difference between XY SDpop model and NULL model calculated over 10 kb sliding windows. An elevated BIC difference (>2000) corresponds to more support for the XY model (XX-XY sex determining system) compared with the NULL model (no differentiation between males and females) for a particular 10 kb window. **C)** Male-female genome-wide *F*_ST_ for linkage group 13 (scaffold12). **D)** BIC differences between XY and NULL SDpop model for linkage group 13 (scaffold12).

To further investigate sex-specific genomic differentiation within this region, we compared sequencing coverage between males and females. Because the reference genome was derived from a male individual, reduced mapping of female reads is expected in Y-linked or diverged XY regions. Consistent with this expectation, males showed higher sequencing coverage than females across the candidate SD region, whereas females exhibited locally reduced coverage (Fig. 4A–B). This pattern suggests the incorporation of Y-linked sequence, or of sufficiently diverged XY sequence, into the reference assembly.

**Figure 4:**
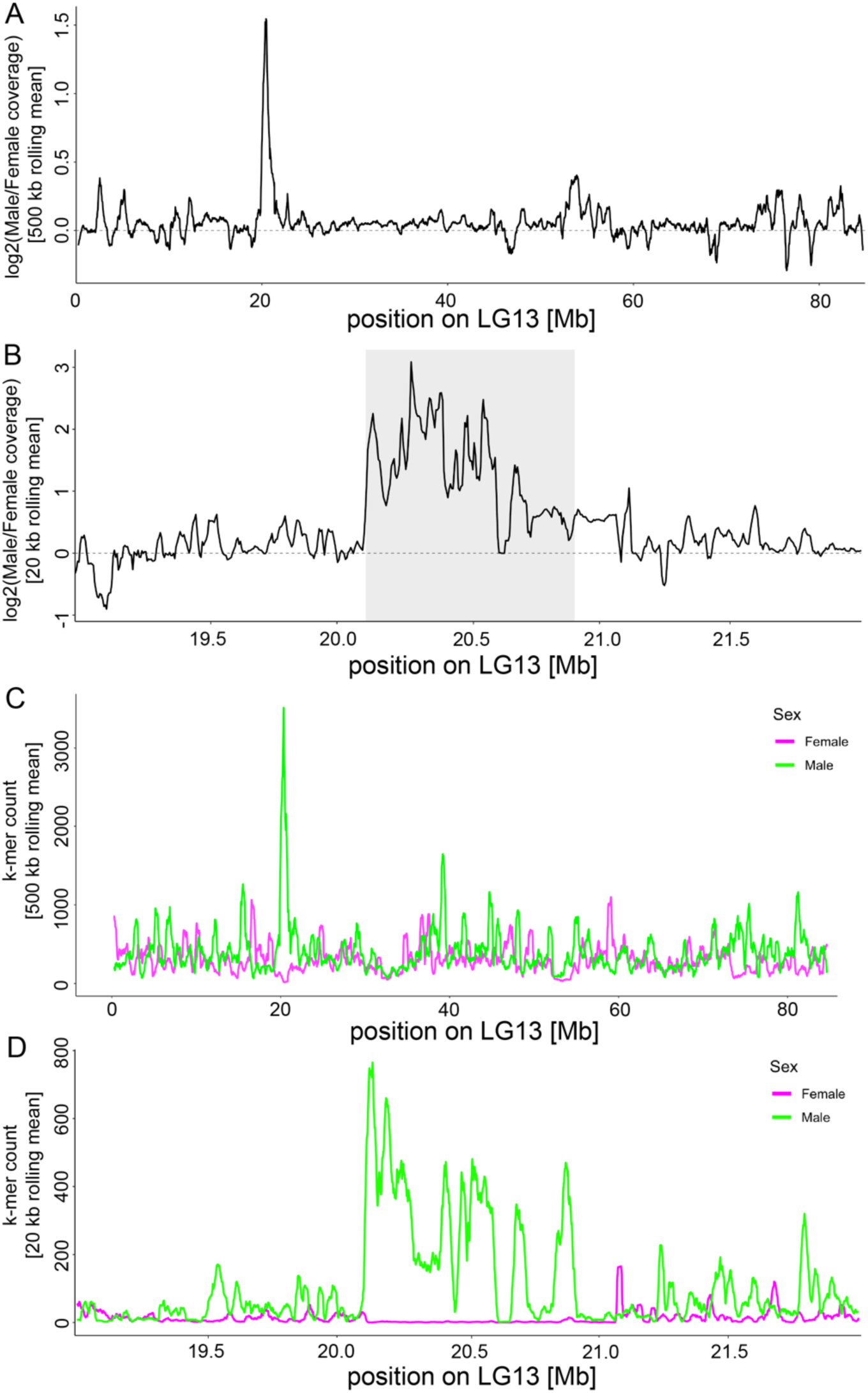
Sex-specific coverage and k-mer patterns across linkage group 13 support a male-specific region in quagga mussels. **A)** Genome-wide profile of the log2 male:female read coverage ratio across linkage group 13 (LG13; scaffold12), calculated from 5 kb bins and smoothed with a 500 kb rolling mean using 19 males and 19 females. Values near zero indicate similar coverage between sexes, whereas positive values indicate higher male coverage. Most of LG13 shows similar coverage between sexes, except for a localized male-biased region around ∼20–21 Mb. **B)** Zoomed-in view of the 19–22 Mb interval on LG13 showing the log2 male:female read coverage ratio calculated from 5 kb bins and smoothed with a 20 kb rolling mean. The shaded region indicates the candidate sex-determining (SD) region. Coverage is strongly male-biased across much of this interval, consistent with the presence of Y-linked sequence in the male-derived reference genome. **C)** Genome-wide distribution of male-specific and female-specific k-mer counts (500 kb rolling mean) along LG13. Male-specific k-mers are strongly enriched in the SD region, whereas female-specific k-mers remain at background levels. **D)** High-resolution view (20 kb rolling mean) of male-specific and female-specific k-mer counts across the 19–22 Mb interval. A pronounced cluster of male-specific k-mers delineates an ∼800 kb male-specific region, consistent with the presence of Y-linked sequence absent from females.

To independently validate the presence of a localized XY SD region, we performed a SNP calling-free k-mer analysis. Sex-specific 31-bp k-mers were identified in 19 males and 19 females using two abundance thresholds (minimum counts of 20 and 40; Table S3) and subsequently mapped to the reference genome. Because the more stringent dataset (minimum count of 40) yielded higher mapping rates, it was retained for downstream analyses (Table S3). Genome-wide, male- and female-specific k-mers mapped relatively uniformly across chromosomes, with slightly higher coverage in males, consistent with the male-derived reference genome (Table S4). In stark contrast, within the SD region on LG13, male-specific k-mer counts were much higher than female-specific k-mers (Fig. 4C). A detailed investigation of the SD region showed that female-specific k-mers were almost entirely absent, resulting in near-zero counts across most of the region (Fig. 4D). This striking asymmetry provides independent evidence for a localized Y-linked SD region in quagga mussels.

### Gene content of the quagga mussel SD region identifies FoxL2-Y, a FoxL2-derived locus

Inspection of gene annotations within the ∼800 kb SD region on LG13 revealed 41 predicted gene models (Table S5). Many of these encoded C-type lectin proteins, a gene family characterized by rapid evolution and frequent involvement in reproductive and immune-related processes. Among the annotated genes, one locus (jg4150) was annotated as *FoxL2*. Because *FoxL2* is a deeply conserved female-associated gene across metazoans, its presence within the LG13 candidate XY SD region warranted further examination. To place this locus in a genome-wide context, we conducted a homology search of the quagga genome using the zebra mussel *FoxL2* sequence as a query. In addition to the LG13 SD locus, this search identified three autosomal *FoxL2*-like paralogs clustered on LG8 (jg24628.t1, jg24629.t1, jg24630.t1) and two autosomal *FoxL2*-like paralogs on LG14 (jg40939.t1 and jg41002.t1). Among these, jg24628, jg24630.t1, and jg40939.t1 each have complete forkhead domains and show high similarity to zebra mussel *FoxL2* (90%, 89%, and 88% identity respectively in full length alignments). Inferred amino acid sequences for jg24629.t1 and jg41002 show partial forkhead domains, consistent with structural divergence.

Targeted sequence comparisons between the LG13 SD locus and the *FoxL2* genes on LG8 revealed that the SD-associated locus represents a duplicated and structurally divergent paralog of *FoxL2* (jg24630.t1); this locus is hereafter referred to as *FoxL2-Y* (Fig. 5A; Table S6). Exon–intron structure analysis showed that *FoxL2-Y* retains overall exon organization and extensive coding-sequence homology with *FoxL2* (jg24630.t1) (Fig. 5A; Table S6). However, *FoxL2-Y* differs from *FoxL2* by the presence of a large insertion composed mostly of simple repeats within the first intron, resulting in marked intron expansion and disruption of collinearity (Fig. 5A). Within the exonic regions, *FoxL2-Y* carries multiple nucleotide substitutions and short indels, including a six–amino-acid deletion near the 5ʹ end and several nonsynonymous substitutions (Figs. S5-S6). Notably, *FoxL2-Y* also contains a premature stop codon within the forkhead (FH) domain (Table S6). Amino acid alignment highlights truncation of the FH domain in *FoxL2-Y* despite overall sequence conservation across the remainder of the coding region (Fig. S6). Together, these features identify *FoxL2-Y* as a *FoxL2*-derived paralog with preserved gene structure but a disrupted FH domain and substantial divergence in predicted coding potential.

**Figure 5:**
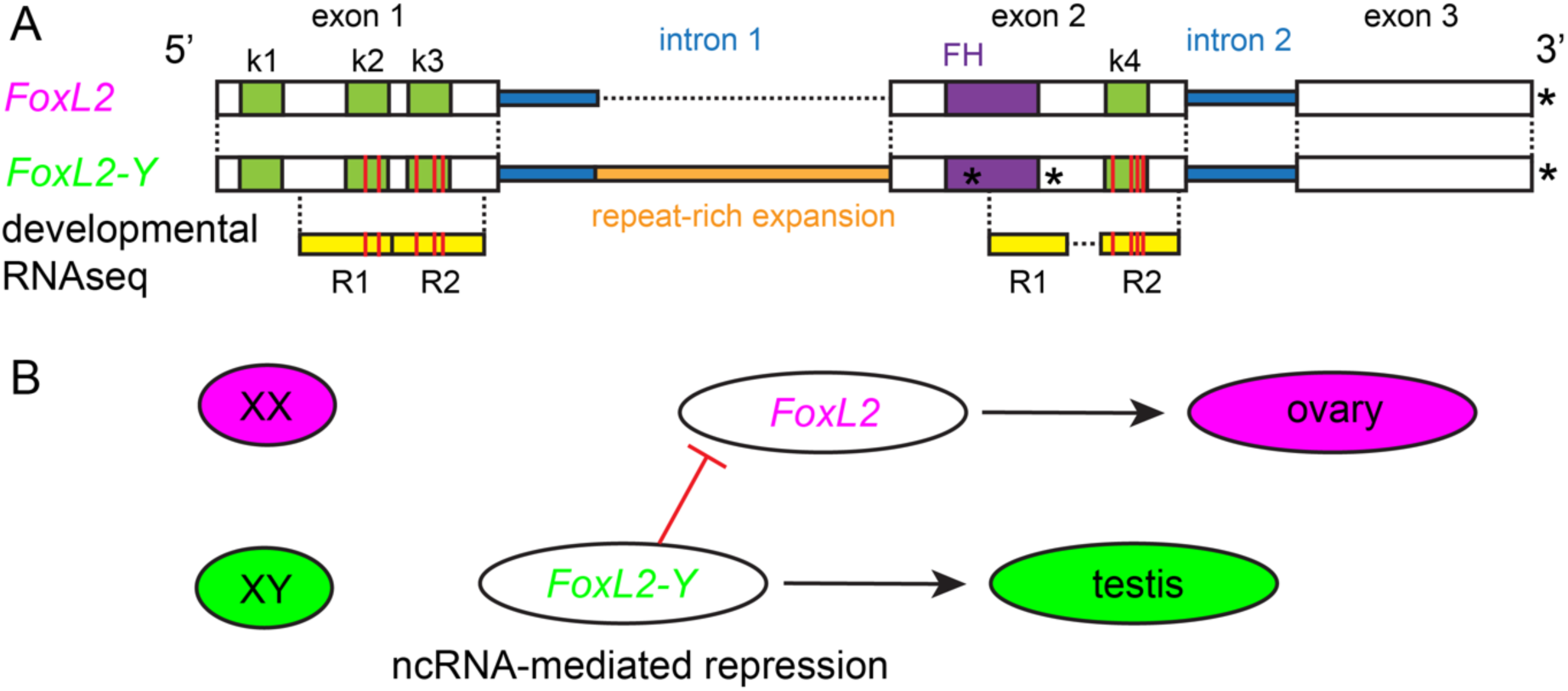
Structural divergence and putative regulatory role of the Y-linked *FoxL2* paralog (*FoxL2-Y*) in quagga mussels. **A)** Schematic comparison of the canonical *FoxL2* gene (jg24630.t1) and the Y-linked paralog *FoxL2-Y* identified within the quagga mussel sex-determining region. Exons are shown as boxes and introns as connecting lines, with exon and intron numbering indicated above. The conserved forkhead (FH) domain is highlighted in purple. *FoxL2-Y* retains overall exon structure and coding-sequence homology with *FoxL2* but contains a large repeat insertion within the first intron and multiple disruptive mutations, including a premature stop codon (asterisk) within the FH domain. Positions of diagnostic *FoxL2-Y*–specific k-mers (k1–k4; dark green) used for reference-free detection are indicated. Two embryonic RNA-seq paired-end reads uniquely matching *FoxL2-Y* are shown beneath the gene model (diagnostic variants in red), confirming rare transcription during early development. Full nucleotide and amino acid alignments are provided in Figs. S5 and S6. **B)** Conceptual model for sex determination in quagga mussels. In XX individuals, FoxL2 activity promotes ovarian development. In XY individuals, early, transient expression of *FoxL2-Y* is hypothesized to repress FoxL2 through a noncoding RNA–mediated mechanism, thereby biasing development toward the male pathway. This model is consistent with the genomic localization, structural features, and stage-restricted transcription of *FoxL2-Y* but remains to be functionally tested.

To place the quagga SD-region locus in a broader comparative context, we examined *Fox*, *Sox*, and *DMRT* gene-family composition and phylogenetic relationships across *Dreissena* and other bivalves (Figs. S7–S10). Gene copy-number comparisons indicated broadly conserved *Sox* and *DMRT* repertoires across taxa and confirmed the absence of a *DMRT-1L* ortholog in both zebra and quagga mussels (Fig. S7). The Fox phylogeny supported assignment of the LG13 SD-region copy to the FoxL2 lineage, consistent with *FoxL2-Y* representing a derived paralog of canonical *FoxL2* rather than a member of another Fox subfamily (Fig. S8). In contrast, the *Sox* and *DMRT* phylogenies did not reveal an obvious quagga-specific candidate comparable to *FoxL2-Y* and mainly served to confirm expected gene-family composition in both *Dreissena* species (Figs. S9 and S10).

### K-mer–based transcriptomic screening reveals rare and development-restricted FoxL2-Y expression

Because *FoxL2-Y* and other *FoxL2* paralogs on LG8 and LG14 share extensive sequence similarity across much of their coding regions, conventional read-mapping approaches cannot reliably distinguish transcripts derived from the two loci. We therefore used a reference-independent strategy based on diagnostic k-mers spanning *FoxL2-Y*–specific indels and/or linked nucleotide substitutions to screen transcriptomic datasets. We queried 23 publicly available embryonic and developmental RNA-seq libraries spanning six developmental stages, from unfertilized eggs to juvenile stages, as well as newly generated adult tissue-specific RNA-seq libraries. *FoxL2-Y*–specific k-mers were detected at extremely low abundance in two independent early developmental libraries: one trochophore-stage library and one early veliger-stage library (Table S7). Extraction and reconstruction of *FoxL2-Y*–positive paired-end reads confirmed the presence of multiple linked *FoxL2-Y*–specific diagnostic variants within individual transcript fragments, including variants located downstream of the premature stop codon, validating their locus-specific origin and indicating transcription across multiple exons of *FoxL2-Y* (Fig. 5A). No *FoxL2-Y*–specific k-mers were detected in earlier embryonic stages, later larval stages, juvenile samples, or adult tissues, including testes (Table S7; Table S8). In contrast, *FoxL2*-specific k-mers were detected at similar or later developmental stages in quagga mussels and in mixed-species libraries containing zebra mussel larvae (Table S7). In adult quagga mussels, *FoxL2* showed ovary-biased expression relative to other tissues (Table S8).

### Conceptual model of sex determination in quagga mussels

Based on the male-specific genomic localization of *FoxL2-Y*, its derivation from the conserved female-associated gene *FoxL2*, the presence of a premature stop codon within the FH domain, and its rare transcription during early development, we propose a conceptual model summarizing the observed patterns of sex determination in quagga mussels (Fig. 5B). In this model, *FoxL2* activity is associated with female development, whereas the Y-linked *FoxL2-Y* locus is hypothesized to influence male development through an early, transient regulatory effect, potentially independent of canonical protein-mediated transcriptional activity. This model integrates the contrasting genomic and expression profiles of *FoxL2* and *FoxL2-Y* and is intended as a descriptive framework to guide future functional investigations rather than as a demonstration of causality.

## Discussion

### Contrasting and rapidly diverging genomic architectures of sex determination in quagga and zebra mussels

In this study, we investigated the genomic architecture of sex determination in two closely-related invasive freshwater mussels using chromosome-scale reference genomes and whole-genome resequencing data. A key outcome is the generation of a chromosome-scale reference genome for the quagga mussel, which provides a novel resource for molluscan genomics and enables fine-scale analyses of sex-linked regions that were previously not possible in this species. In addition, we leveraged whole-genome population resequencing to identify sex-linked genomic signatures, complementing previous bivalve studies that have primarily relied on comparative genomics (*23*) or transcriptomic approaches (*21*). Together, these genomic resources revealed strikingly different signatures of sex determination between quagga and zebra mussels, which diverged approximately 10 million years ago (*31*). In both species, genome-wide male–female differentiation was negligible and no heteromorphic sex chromosomes were detected, indicating that sex determination is associated with localized genomic differences rather than large-scale chromosomal divergence. Despite these shared features, the underlying genomic architectures differed markedly. In zebra mussels, multiple ZW genomic regions distributed across at least two linkage groups exhibited consistent male–female differentiation, whereas in quagga mussels all XY sex-linked signals converged on a single, sharply delimited genomic region. These contrasting patterns point to fundamentally different sex-determining architectures despite the relatively recent evolutionary separation of the two species, and underscore the evolutionary lability of sex determination systems in dreissenid mussels.

While the absence of an appropriate outgroup prevents direct inference of the ancestral state, the identification of *FoxL2-Y* as a derived paralog of the conserved female-associated gene *FoxL2* suggests that the localized XX–XY system in quagga mussels may represent a derived condition, whereas the polygenic architecture observed in zebra mussels may more closely reflect the ancestral state. Comparable patterns of rapid turnover in sex-determining mechanisms have been documented across diverse animal taxa, including fishes, amphibians, and reptiles, where closely-related species can differ in heterogametic sex and in the genetic basis of sex determination (*33*, *39*, *42*–*44*). Such transitions can occur without the evolution of strongly differentiated sex chromosomes, by involving subtle genomic changes and shifts in linkage relationships. The dreissenid mussels thus add to growing evidence that sex determination can evolve rapidly through changes in genomic architecture rather than through gradual chromosomal differentiation.

### Genomic evidence supports a polygenic ZZ–ZW sex determination architecture in zebra mussels

In zebra mussels, multiple independent lines of genomic evidence support a polygenic ZZ–ZW sex determination architecture. We identified eight candidate sex-linked regions distributed across three linkage groups, each characterized by male–female *F*_ST_ values close to the theoretical expectation for sex-linked loci and enrichment for ZW-type SNPs. Four of these regions across two linkage groups additionally overlapped with candidate windows identified by model-based SDpop analyses, providing convergent support across complementary approaches. Together, these patterns are consistent with a system in which sex determination is governed by multiple loci located on different linkage groups rather than by a single master locus (*14*). The three candidate regions on LG5 are particularly notable because each encompasses a single annotated gene. One of these single-gene regions contains *FoxL2*, a gene with a well-established role in ovarian development and female sex determination across bivalves (*21*, *23*) and metazoans more broadly (*45*, *46*), making it a compelling candidate in a ZW context. In zebra mussels *FoxL2* exists a single-copy gene, in contrast to the duplicated paralogs identified in quagga mussels. The two other single-gene regions harbor a tudor domain–containing protein and a homolog of nucleolar complex protein 2, which is involved in negative regulation of RNA polymerase II–mediated transcription. Although these genes have not been directly implicated in sex determination in bivalves, their regulatory functions are consistent with roles in developmental decision-making, including sex determination (*47*–*50*).

Additional candidate regions on LG9 and LG13 include genes annotated as transcription factors, a functional class frequently associated with sex-determining pathways across animals, including canonical SD genes such as *FoxL2* and *DMRT-1* (*51*, *52*). Notably, several candidate regions on LG13 harbor long non-coding RNAs, a gene class increasingly recognized for regulatory roles in developmental processes, including sex determination (*53*–*57*). The recurrence of regulatory genes and non-coding elements among candidate regions therefore supports a model in which sex determination in zebra mussels arises from the combined effects of multiple regulatory loci (*14*). Although the genomic signatures observed here are consistent with a polygenic architecture, they do not exclude the possibility that one or a few dosage-sensitive regulatory genes—such as *FoxL2*—may occupy a central position within this network. In such a scenario, variation at multiple loci could converge on the regulation or effective dosage of *FoxL2*, generating a polygenic genomic signal without implying equivalent functional contributions of all loci. Polygenic sex determination remains relatively rare among well-characterized systems, but has been documented mostly in fishes, as well as in reptiles, amphibians, reptiles, mammals, houseflies and copepods, where sex is determined by the additive or epistatic effects of multiple loci distributed across the genome (*14*). Although functional validation is currently lacking, the consistency of genomic signals across multiple regions, chromosomes, and analytical approaches strongly suggests that sex determination in zebra mussels is genuinely polygenic rather than the result of a single cryptic sex-determining locus.

### A localized XX–XY genomic region and a FoxL2-derived candidate locus in quagga mussels

In contrast to the diffuse, polygenic pattern observed in zebra mussels, quagga mussels exhibited a highly localized genomic signature of sex determination consistent with an XX–XY system. A single ∼800 kb region on LG13 showed elevated male–female differentiation, a strong SDpop signal, reduced sequencing coverage in females, and a pronounced enrichment of male-specific k-mers, providing independent genomic evidence for a localized Y-linked region. Within this region, we identified *FoxL2-Y*, a divergent paralog of the canonical FoxL2 gene. Detailed sequence analyses revealed that *FoxL2-Y* retains extensive nucleotide and amino-acid sequence similarity with *FoxL2* across all annotated exons, despite the accumulation of multiple nonsynonymous substitutions and short deletions. This overall conservation contrasts with a marked structural disruption within the coding sequence, including a premature stop codon located within the forkhead (FH) domain, which is intact in canonical FoxL2. As a result, *FoxL2-Y* lacks an uninterrupted FH domain and is unlikely to encode a fully functional DNA-binding transcription factor, despite preserving substantial homology across the remainder of the coding region. Transcriptomic screening using diagnostic k-mers detected *FoxL2-Y* expression at extremely low abundance and restricted to early development, with *FoxL2-Y*–specific reads identified in two independent developmental libraries corresponding to the trochophore and early veliger stages. Notably, one read pair mapped downstream of the premature stop codon within exon 2, indicating that transcription extends beyond the predicted truncation point. Although these observations do not establish a causal role for *FoxL2-Y*, they are consistent with *FoxL2-Y* being transcriptionally active during a narrow developmental window and tightly linked to the sex-determining region in quagga mussels.

In many metazoans, including fruit flies, mammals, fishes, and amphibians, male sex determination is commonly associated with genes of the *DMRT* family, which have repeatedly been recruited as key male-determining factors across distant lineages (*52*, *58*). In bivalves, a bivalve-specific paralog of *DMRT-1*, *DMRT-1L*, has recently been proposed as a conserved male-determining gene based on comparative genomic and evolutionary analyses (*23*). Under this framework, one might expect species with an XX–XY system to rely on *DMRT-1L*, as has been suggested for the invasive golden mussel *Limnoperna fortunei* (*59*). However, recent analyses of high-quality bivalve genomes indicate that DMRT-1L has been lost in the order Myida (which includes *Dreissena*), and potentially across the broader superorder Imparidentia, although inference is complicated in some taxa by reliance on transcriptomic data (*23*, *60*). The absence of *DMRT-1L* in quagga mussels implies that, if sex determination is male-heterogametic, it must be mediated by an alternative locus. In this context, the emergence of a *FoxL2*-derived locus within the quagga mussel SD region represents an intriguing and parsimonious alternative. *FoxL2* is a deeply conserved female-associated gene across metazoans, and duplication followed by male-specific genomic linkage could provide a route to the evolution of a novel male-determining regulatory element. While functional validation will be required to resolve the precise role of *FoxL2-Y*, its genomic localization, retained exon-wide homology to *FoxL2*, structural disruption of the FH domain, and early developmental expression pattern make it a compelling candidate locus in a lineage that has lost the canonical bivalve male-determining gene.

Although rare, duplication-driven rewiring of core sex-pathway genes has been previously documented. In *Xenopus laevis*, the W-linked female-determining gene DM-W arose through duplication of the ancestral male-associated gene *DMRT-1* and functions as the dominant trigger of ovary development in a derived ZW system (*61*, *62*). Phylogenomic analyses indicate that this system originated following allotetraploidization in the *X. laevis* lineage approximately 17–18 million years ago (*63*), demonstrating that novel sex-determining loci can emerge through duplication of conserved pathway genes on relatively recent evolutionary timescales. In light of this precedent, the hypothesis that a *FoxL2*-derived duplicate contributes to male determination in quagga mussels is evolutionarily plausible and consistent with known mechanisms of SD system turnover.

### Possible modes of action of the FoxL2-Y locus

Although the genomic localization and developmental expression pattern of *FoxL2-Y* are consistent with a role in sex determination, its precise mode of action remains unknown. The presence of a premature stop codon within the FH domain makes it unlikely that *FoxL2-Y* functions as a conventional DNA-binding transcription factor. Importantly, however, the preservation of exon structure and the detection of transcripts extending beyond the truncation point suggest that *FoxL2-Y* is not a transcriptionally inert pseudogene. Together, these features raise the possibility that *FoxL2-Y* may act through a non-canonical regulatory mechanism rather than as a protein-coding gene. Increasing evidence across metazoans indicates that long non-coding RNAs (lncRNAs) and other regulatory RNAs can play central roles in developmental decision-making, including sex determination, by modulating chromatin state, transcriptional activity, or RNA stability of key pathway genes (*53*–*56*, *64*, *65*). One plausible hypothesis is that *FoxL2-Y* functions as a regulatory RNA that interferes with *FoxL2* expression or activity during a narrow early developmental window. Such interference could occur through (i) competition for shared regulatory factors, (ii) recruitment of chromatin-modifying complexes, or (iii) RNA–RNA interactions, even in the absence of protein-coding capacity (*53*, *66*, *67*). Among these possibilities, competition for shared regulatory factors may be particularly plausible given the extensive sequence similarity retained between *FoxL2-Y* and *FoxL2* across their coding exons, which could allow *FoxL2-Y* transcripts to engage molecular partners normally involved in *FoxL2* regulation without requiring an intact DNA-binding domain (*53*). Notably, several candidate sex-linked regions in zebra mussels also contain putative long non-coding RNAs, raising the possibility that regulatory RNA–mediated mechanisms could be recurrent features of sex determination in *Dreissena*. We emphasize that these hypotheses remain speculative and require functional validation. Nevertheless, the emergence of a *FoxL2*-derived, structurally modified locus that is male-specific, transcriptionally active during early development, and embedded within a localized SD region is consistent with models in which duplication and regulatory rewiring of conserved sex pathway genes generate novel sex-determining elements (*61*). *FoxL2-Y* thus represents a promising candidate for further investigation into non-canonical and regulatory modes of sex determination in quagga mussels.

### Genome dynamism and haploid selection as putative mechanisms underlying sex determination turnover

The genomic context in which sex determination systems evolve likely plays an important role in facilitating transitions between alternative architectures. Comparative analyses revealed extensive chromosomal rearrangements between quagga and zebra mussels despite similar genome sizes and identical chromosome numbers, indicating a highly dynamic genomic landscape in this clade. Structural rearrangements, gene duplications, and transposable-element activity can reshape recombination landscapes and generate novel linkage relationships, thereby creating opportunities for the origin and spread of new sex-linked regions (*68*, *69*). Such genome dynamism has been associated with rapid turnover of sex-determination systems in other taxa, particularly fishes and amphibians, where closely related species often differ in both sex-determining loci and heterogametic sex (*70*, *71*). In this framework, duplication and genomic relocation of regulatory genes—such as the FoxL2-derived locus identified here—can provide raw material for the evolution of novel sex-determining elements when embedded in permissive genomic contexts.

Within a dynamic genomic background, selection acting at the haploid stage may further promote rapid sex determination turnover (*72*, *73*). In quagga mussels, the sex-determining region is notably enriched in *C-type lectin* genes and *peptide G protein-coupled receptors*, gene families notably implicated in innate immune recognition and reproductive signaling, respectively (*74*, *75*). Carbohydrate–lectin interactions are well established as key mediators of sperm–egg recognition across metazoans (*76*–*78*), and lectin-mediated binding has been documented in bivalves, including *Dreissena* species, where characteristic carbohydrate patterns have been detected on both eggs and sperm (*79*, *80*). In externally fertilizing species such as quagga and zebra mussels, selection acting during the gametic stage—through gamete competition, compatibility, or fertilization efficiency—has been proposed as a powerful force shaping reproductive traits (*81*, *82*).

Theoretical work has shown that haploid selection can favor the spread of new sex-determining alleles, particularly when these arise in genomic proximity to loci under haploid selection, although strict physical linkage is not always required (*72*, *73*). Under a “linkage-strengthening” scenario, haploid selection may accelerate the stabilization of a newly emerged sex-determining locus embedded within a region enriched for gamete-interaction genes, thereby facilitating transitions between ZZ–ZW and XX–XY systems over relatively short evolutionary timescales. While direct evidence for haploid selection in dreissenid mussels is currently lacking, several observations in quagga mussels are consistent with this model: the presence of a single localized Y-linked SD region, the emergence of a *FoxL2*-derived male-specific candidate locus, and the clustering of genes plausibly involved in gamete recognition within the same genomic block. In contrast, the polygenic ZW architecture observed in zebra mussels may reflect a different historical equilibrium, consistent with theoretical predictions that haploid selection can promote frequent transitions between alternative sex determination systems. More generally, theoretical models predict that polygenic sex determination is intrinsically unstable in the presence of sex-specific selection, which favors the evolution of a single dominant sex-determining locus linked to sexually antagonistic alleles (*83*).

### Broader implications for invasion genomics and the development of genetic biocontrol

Beyond their relevance to sex determination, our results highlight dreissenid mussels as a promising system for studying how genome dynamism and high levels of standing genetic variation may facilitate rapid evolutionary responses in invasive species (*84*, *85*). The extensive chromosomal rearrangements observed between quagga and zebra mussels, together with their high levels of genomic variation and heterozygosity, are consistent with a genomic context that may promote rapid adaptation from standing variation following invasion. While these features cannot be directly linked to invasion success based on the present data, they underscore the evolutionary flexibility of these species and provide important context for understanding their rapid spread across novel environments.

From an applied perspective, resolving the genomic basis of sex determination is directly relevant to emerging genetic biocontrol strategies that aim to manipulate reproduction or sex ratios in invasive populations (*8*, *9*). Our results suggest that the feasibility of sex-ratio–based biocontrol may strongly depend on the underlying SD architecture. In zebra mussels, the polygenic nature of sex determination suggests that targeting sex determination through genome editing would be complex and likely less tractable than in systems with a single master locus. Whether sex-ratio distortion could nonetheless be feasible will depend on whether cumulative dosage effects across multiple loci converge on a small number of key regulators, such as FoxL2, a hypothesis that remains to be explicitly tested. In contrast, the discovery of a single, localized XX–XY sex-determining region in quagga mussels, together with a *FoxL2*-derived candidate locus, suggests that sex-ratio distortion approaches could be conceptually more straightforward in this species.

The identification of *FoxL2-Y* as a putative male-determining locus on a homomorphic sex chromosome in quagga mussels is analogous to the case in disease vector mosquitos of the genus *Aedes*. Like zebra and quagga mussels, *Aedes* mosquitos are globally invasive (*86*). In *A. aegypti* the gene *Nix* has been identified as the “M factor” male determinant on homomorphic sex-determining factor (*87*). *Nix* has been shown to be required for male sex determination and sufficient to convert genetic females into fertile males in both *A. aegypti* and *A. albopictus* (*88*–*90*). The identification of *Nix* and downstream effectors in *Aedes* species has opened the door to novel approaches for genetic biocontrol through engineering of sex determination and other reproductive traits (*91*–*93*). The identification of *FoxL2-Y* in quagga mussel presents a similar opportunity to pursue development of genetic biocontrol targeting sex ratios or reproductive output. Despite this potential, we emphasize that such applications remain currently highly speculative. Functional validation of *FoxL2-Y*, the establishment of reliable laboratory reproduction, and the development of efficient genome-editing and delivery methods in early quagga embryos all represent substantial technical challenges (*9*). In addition, any consideration of gene drive–based interventions must carefully address regulatory, ecological, and ethical constraints associated with the release of self-propagating genetic constructs into natural ecosystems (*94*, *95*).

More broadly, the contrasting sex-determining architectures uncovered here illustrate how closely related species can arrive at fundamentally different genetic solutions to the same developmental problem over relatively short evolutionary timescales. By integrating chromosome-scale genomics, population resequencing, and transcriptomic screening, this study provides a framework for disentangling sex determination in non-model invasive species. Future work combining functional assays, expanded population sampling across native and invaded ranges, and detailed analyses of gamete biology will be essential to resolve causative mechanisms and to understand how genomic architecture, selection, and life history interact to shape the evolution of sex determination in ecologically and economically impactful species.

## Methods

### Quagga mussel reference genome

#### Sampling and DNA extraction

Adult quagga mussels for genomic DNA sequencing were collected from the forebay of Davis Dam, near Bullhead City, Arizona, USA (35.196077, -114.56899) on May 02, 2018. Adult mussels were collected and maintained in aerated fresh water until they were sacrificed. Individuals were dissected and sexed under a stereo microscope with sterile dishes, forceps, and scalpels used for each individual. Tissue samples including foot, gills, and gonads were placed in individual screw-top cryotubes. Samples of male individuals were flash frozen in liquid nitrogen and stored in a -80°C freezer. DNA extraction and isolation was performed with the Qiagen Blood & Cell Culture Kit (QIAGEN Inc., USA) using Genomic Tip 100/G. Following Proteinase K digestion, the samples were pre-filtered through a nylon screen filter basket with a 100 μm pore size (Corning Sterile Cell Strainer – manufacturer product number 431752) prior to binding the sample to the Genomic Tip filter. DNA yield was measured with a Qubit 4 fluorometer (Thermo Fisher Scientific, USA) using the Qubit dsDNA BR Assay Kit to label samples. DNA purity was determined by measurement of light absorption values at 260 nm and 280 nm using a Cary-60 spectrophotometer with a 1.0 mm microcell cap for measurement of small volumes.

#### Illumina and PacBio sequencing

Two technologies, Pacific Biosciences (PacBio) Sequel Single Molecule Real Time (SMRT) sequencing and Illumina HiSeq paired-end (PE) sequencing, were used to collect genomic data for the quagga mussel. A goal of 100x coverage from both sequencing technologies was chosen as a target that would maximize the data available to develop a high-quality assembly. No direct measure of the quagga mussel genome size was available, however, an estimate of 1,660 Mbp was available for the zebra mussel (*Dreissena polymorpha*) (*96*). This was considered an appropriate estimate and was used for experimental design. To achieve 100x coverage, sequencing was targeted for an output of 160,000 Mbp from both PacBio Sequel SMRT and Illumina HiSeq PE technologies. Genomic DNA extracted from a male mussel labeled ‘Drb016’ was sent to GENEWIZ for PacBio Sequel SMRT and Illumina HiSeq PE sequencing. GENEWIZ generated 664,901,022 Illumina PE reads and 11,199,225 PacBio Sequel Continuous Long Reads (CLR) totaling 165,845 Mbp. CCS reads from the PacBio Sequel runs had a mean read score of 0.988, with an average of 9.8 read passes per insert. In total, there were 33,350 CCS reads totaling 308 Gbp.

#### Genome size and heterozygosity estimation

GenomeScope (Vurture et al. 2017) was used to estimate the quagga mussel genome size and heterozygosity based on the Illumina PE reads. First, the raw reads were quality and adapter trimmed and filtered using Trimmomatic (*97*) with the following options: “ILLUMINACLIP:TruSeq3-PE-2.fa:2:30:10 LEADING:10 TRAILING:10 SLIDINGWINDOW:4:15 MINLEN:50” where TruSeq3-PE-2.fa is a FASTA file containing the sequences of the Illumina TruSeq sequencing adapters. Jellyfish 2.2.10 (https://github.com/gmarcais/Jellyfish) was then used to count k-mers in the quality and adapter trimmed/filtered data and generate a histogram for GenomeScope. The options for jellyfish count used were “-C -m 21 -s 19000000000.” The GenomeScope web server (accessed April 7, 2020) was then used to calculate the GenomeScope profile and estimate the genome length and heterozygosity.

#### Genome assembly

PacBio reads including CCS reads were converted from BAM to FASTA format with bam2fasta (https://github.com/PacificBiosciences/bam2fastx). Reads were then assembled using Canu 1.8 (*98*) with the following options: “-genomeSize=2g -pacbio-raw” where 2g indicates that the estimated genome size for the quagga mussel was on the order of 2 Gbp. Given the high heterozygosity of the quagga mussel genome (see Results), redundant haplotigs retained in the primary Canu assembly were removed using purge_dups 1.0.1 (*99*). A config.json file was prepared to request 16 cores and 50 Gb of RAM on the HPC cluster of the University of Alabama. Otherwise the default settings were used. The resulting ‘purged’ genome assembly was then ‘polished’ to correct errors that may have been introduced into the assembly by low-quality PacBio subreads using the version of POLCA bundled with MaSuRCA 3.3.5 (*100*). The Trimmomatic-trimmed Illumina PE reads described above were used.

#### Hi-C scaffolding and genome quality control

The resulting assembly was then scaffolded to chromosome-level using Phase Genomics Hi-C. Intact frozen tissue was sent to Phase Genomics for library preparation and sequencing. Ideally, tissue for Hi-C scaffolding would come from the same individual used from primary sequencing and contig assembly. However, all available tissue from the mussel Drb016 was used for DNA extraction to accomplish PacBio Sequel and Illumina HiSeq. Therefore, tissue from a second male individual was used for Hi-C library preparation. Phase Genomics produced 119,158,848 Illumina reads from the constructed Hi-C library. At the time where the Hi-C scaffolding was conducted, there was no accurate quagga mussel karyotype available. Therefore, given that the zebra mussel genome consists of 16 pairs of chromosomes (*26*, *101*), the initial Hi-C scaffolding with the Phase Genomics Proximo pipeline was constrained to be between 14 and 20. Karyotype information obtained a posteriori confirmed that quagga mussels also have 16 pairs of chromosomes (see below). Contiguity and completeness of the genome assembly were assessed at each step with QUAST 5.0.2 (*102*) and BUSCO 4.0.6 (*103*), respectively.

#### Transcriptomics for genome annotation

Adult quagga mussels for RNA sequencing were collected from the forebay of Davis Dam, near Bullhead City, Arizona, USA (35.196077, -114.56899) on May 02, 2018. Six samples were prepared for transcriptome sequencing (i.e., RNAseq) for genome annotation. These included two whole mussels and four individual tissue samples (gill, foot, ovary, and testes + digestive gland combined). RNA isolation and purification were performed with the RNAEasy Mini kit (QIAGEN Inc., USA), following the manufacturer’s protocol, including DNAse treatment. RNA yield was quantified using the Qubit RNA BR Assay Kit (Thermo Fisher, USA) and RNA purity was assessed by spectrophotometric measurement of the 260/280 ratio on a Cary-60 (Agilent, USA). Individual library preparation was performed with the Illumina TruSeq Stranded mRNA kit and libraries were run with Illumina 150 bp PE sequencing with a target depth of 200 million paired-end reads per library. After sequencing, between 198.6 million and 255.7 million paired-end reads per library were recovered. In all samples the percentage of bases with a quality score equal to a greater than 30 was 90% or higher. Each transcriptome was assembled in Trinity 2.10.0 using the built-in quality and adapter trimming/filtering and digital normalization options with the default settings. A combined assembly containing all reads from all transcriptomes was also produced. Assembled transcriptomes were translated with TransDecoder 5.5.0 (*104*) using the UniProt SwissProt database (The Uniprot Consortium, 2014) and PFAM (Pfam-A.hmm) version 27 (*105*) to guide translation.

#### Genome annotation

For genome annotation, we identified and masked repetitive DNA and then used both RNA-seq data and protein sequence data as evidence for protein-coding gene prediction. Repeats in the Hi-C scaffolds were annotated with RepeatModeler 2.0.1 with the --LTRStruct option and softmasked with RepeatMasker 4.0.7 with the --gccalc option. The engine used for both programs was rmblast. We ran TrimGalore (*106*) on the newly generated transcriptome reads with the following settings: “-q 30 --illumina --trim-n --length 50.” The resulting quality- and adapter-trimmed and filtered transcriptome reads were then mapped to the repeat masked Hi-C scaffolds using STAR 2.4.0k (*107*) with the --genomeChrBinNbits 15 and --chimSegmentMin 50 options. Our newly sequenced quagga mussel transcriptomes were assembled and translated and described above. These plus select other mollusc proteomes obtained from similarly processed, publicly available transcriptomes and publicly available genome annotations were then aligned to the repeat-masked Hi-C scaffolds with ProtHint 2.6 (*108*) with an e-value cutoff of 1e-25. Annotation of protein-coding genes was then performed with BRAKER 2.1.6 (*109*) using the output of STAR and ProtHint as evidence with the following settings: “--eptmode --softmasking --crf.”

### Karyotyping

#### Quagga mussel collection

Quagga mussels were collected from Lake Michigan and obtained from Dr. Ashley Elgin, NOAA (Great Lakes Environmental Research Laboratory, Muskegon MI, USA). Mussels were transported to the laboratory (Biomilab) (under a scientific collector’s permit for collection and transport of invasive mussels from the Michigan Department of Natural Resources – Fisheries division) where they were maintained at 15°C in aerated aquaria and fed daily with phytoplankton (Rotigrow Nano, Reed Mariculture Inc, USA).

#### Preparation of metaphase spreads

Metaphase spreads were prepared from day-old quagga mussel embryos based on previously published protocols (*101*, *110*). To this end, adult quagga mussels were induced to spawn by exposure to serotonin creatinine sulfate monohydrate (Sigma-Aldrich, USA) according to established protocols (*111*). Following gamete release, *in vitro* fertilization (IVF) was promptly initiated by mixing eggs with sperm cells in sterile aquarium water supplemented with gentamycin (Sigma-Aldrich, USA) and Penicillin/Streptavidin (Gibco, USA) (SAW). Twenty-four hours after fertilization, embryos were collected and incubated in 60 μL/mL colcemid (KaryoMax™Colcemide™, Gibco, USA) for 2-3 hours at 25-27°C to arrest cells in metaphase. Treated embryos were then gently washed twice in SAW and placed in a 75mM potassium chloride hypotonic solution for 10 minutes at room temperature, followed by 10 minutes on ice before being fixed for 3 x 20 minutes in freshly prepared cold Carnoy solution (3:1 methanol/glacial acetic acid). After fixation, embryos were pelleted by gentle centrifugation, the supernatant was removed, and embryos were re-suspended in 100-200 μL fixative before being dropped onto cold histology slides that were immediately placed on a hot plate for several minutes. The next day, metaphase spreads were stained with the nuclear marker 4ʹ,6-diamidino-2-phenylindole (DAPI) and coverslipped with ProLong™ antifade reagent (Thermo Fisher Scientific, USA). Microscopic images of fluorescent spreads were obtained using a VWR inverted fluorescence microscope equipped with an INFINITY5-5 Teledyne camera and software (Lumenera, USA). Spreads in which chromosomes could not be accurately counted were eliminated from the study. As a result, a total of 69 metaphase spreads were analyzed and counted.

### Population genomics of male and female quagga and zebra mussels

#### Individual sampling, DNA extractions and whole-genome resequencing

Adult zebra mussels were sampled in June 2022 in Lake Greifen, Switzerland (N: 47°21’59.3" E: 8°39’55.6") at 1 m depth. 20 males and 20 females were dissected and sexed by visual inspection of the gonads, then preserved in absolute ethanol. Quagga mussels were sampled in August 2021 in Lake Constance, Switzerland (N: 47°41’43.7" E: 9°11’37.3") at 1 m and 60 m depth. They were kept alive in aquaria for 10 months before they were dissected and sexed by visual inspection of the gonads. 10 males and 10 females from each depth were dissected, for a total of 40 quagga mussels. They were subsequently preserved in absolute ethanol. DNA extractions of a total of 80 mussels were performed using the LGC sbeadex Tissue DNA Purification Kit (Cat. no. NAP41405) following manufacturer’s instructions. Genomic DNA integrity was checked on an agarose gel (1,4%) and DNA concentration was measured using Qubit dsDNA broad range assays. 80 individual whole-genome libraries were constructed using the NEBNext Ultra II FS DNA Library Prep Kit (NEB) using half volumes of reagents compared with the manufacturer’s instructions. Enzymatic shearing duration was individually adjusted according to the integrity of extracted DNA: 9 min - 11 min 30 sec minutes for high-molecular weight DNA; 2 min - 2 min 45 sec minutes for partially-degraded DNA and degraded DNA was not sheared. Genomic libraries were individually barcoded using NEBNext(R) Multiplex Oligos for Illumina(R) (Cat.no. E6609), pooled in equimolar concentrations and sequenced on two lanes of one S4 flowcell using Illumina Novaseq 6000 sequencer (150 bp paired-end) at the Functional Genomics Center Zürich (FGCZ), Switzerland.

#### Whole-genome resequencing data processing

All bioinformatic calculations were performed on the HPC cluster Euler (ETHZ), Switzerland. ChatGPT (GPT v4.0 & 5.2) was used for code debugging (custom bash and R scripts). Over both species, an average of 80.7 million reads per individual were obtained, representing an average coverage of 13.8x (2.5x st. dev.) for zebra mussel individuals and 10x (2x st. dev.) for quagga mussel individuals. The quality of raw reads was visually inspected using FastQC v0.11.9 and illumina adapters were trimmed with FastP 0.23.2. For zebra mussel resequencing data, the publicly available chromosome-scale genome of a single male (UMN_Dpol_1.0; NCBI accession: GCF_020536995.1) was used as reference for short read mapping (McCartney et al, 2022). For quagga mussel resequencing data, the chromosome-scale reference genome generated in this study was used. Both reference genomes were indexed with index command in bwa-mem2 v2.2.1. Whole-genome resequencing data of each individual was mapped to the respective reference genome using bwa-mem2 v2.2.1 with default parameters. PCR duplicates were removed and quality filtered using samtools v1.15.1 (MQ<20). Variant calling was performed using command mpileup of BCFtools v1.15.1. The resulting VCF file was filtered using VCFtools v0.1.16 with the following parameters: --remove-indels --maf 0.05 MISS=0.5, QUAL=30, MIN_DEPTH=6x, MAX_DEPTH=17x, MIN_Allele=2 and MAX_Allele=2. Min and max depth values were chosen based on the distribution of average coverage per site. Three quagga mussel individuals with over 70% missing data were removed (Table S1). A total of 24,499,661 and 42,073,009 SNPs were obtained for zebra and quagga mussels, respectively, and used for downstream analyses. Individual missingness and individual mean depth were calculated on the final filtered VCF files for each species using VCFtools commands --missing-indv and --depth, respectively.

#### Comparative genomics in zebra and quagga mussels

For synteny analysis between species, homologous gene pairs were identified between the protein sequences of zebra and quagga mussels using BLASTP v2.14.0 with an E-value threshold of 1e-5. A total of 75,288 genes in zebra mussels and 83,983 genes in quagga mussels were included in the analysis. Putative synteny and collinearity regions were then detected using MCScan toolkit v1.0 (*112*) and visualized by Circos v0.69-9 (*113*). In total, 45,678 collinear genes were identified.

#### Comparison of male-female genomic differentiation

For each species, the following analyses were conducted separately to identify sex-linked genomic regions: Male-female *F*_ST_ was calculated both on a genome-wide level and for each SNPs using VCFtools v.0.1.16 (Weir & Cockerham estimator) (*114*). The expected *F*_ST_ using this estimator for equal sample size of males and females is 0.5 (*41*). Genome-wide association analysis (GWAS) for sex was conducted using PLINK v.1.9. We then identified candidate sex-linked SNPs using allele frequencies in the following way: XY SNPs: the expectation is that these SNPs are heterozygous in males and homozygous in females. We therefore selected SNPs with allele frequencies in males between 0.45-0.55, and allele frequencies in females between 0-0.05 or between 0.95-1. ZW SNPs: the expectation is the other way around, with females being heterozygous and males homozygous. Thus, allele frequencies in females should range between 0.45-0.55 and allele frequencies in males should range between 0-0.05 or between 0.95-1. We refrained from using the strict allele frequencies of 0.5 and 0/1 to account for potential sequencing errors. Since we did not have a priori of the sex determining system of each species, we identified XY and ZW SNPs in both species. Furthermore, we tested if they were 10 kb regions significantly associated with sex using the model-based approach SDpop (*37*). This program allows for testing the model fit for each one of the following models: no sex chromosome model (i.e. NULL model); XY model; ZW model. The program was run for each species and each model on non-overlapping sliding windows of 10 kb. The difference in Bayesian Information Criterion (BIC) from the XY model vs. NULL model of each 10 kb window (also ZW model vs. NULL model) was plotted against the genomic position of each 10 kb window.

#### Identification of sex-linked regions and candidate SD genes

Candidate sex-linked regions were identified based on the presence of tightly clustered XY or ZW single-nucleotide polymorphisms (SNPs) showing male–female *F*_ST_ values of approximately 0.5. Focusing on the overlap between sex-specific SNP clustering and elevated *F*_ST_ allowed us to narrow down candidate regions while reducing background noise. These regions were compared with candidate regions identified using SDpop, considering 10 kb windows for which the Bayesian Information Criterion (BIC) difference between the XY or ZW model and the null model exceeded 2000. Genes located within candidate sex-linked regions and their putative functions were identified using the available genome annotations (zebra mussel: *Dreissena polymorpha* Annotation Release 100, GCF_020536995.1; quagga mussel: present study). When no electronic functional annotation was available, predicted amino-acid sequences were retrieved and homology-based searches were performed using BLASTP against the NCBI non-redundant protein database. For each gene, the best hit with an assigned gene name was reported together with the corresponding E-value. If no hits with a gene name were obtained, the top-scoring hit was reported.

#### Deep characterization of the quagga mussel SD region

Because sex-linked genomic signals in quagga mussels converged on a single, sharply delimited ∼800 kb region—contrasting with the diffuse polygenic patterns observed in zebra mussels (Results)—we performed additional targeted analyses exclusively in quagga mussels. These analyses included sequence-based characterization of genes within the SD region, sex-specific sequencing coverage comparisons, reference-free identification of sex-linked k-mers, and k-mer–based screening of transcriptomic datasets to assess candidate locus expression across developmental stages and adult tissues. Such fine-scale analyses are informative when sex determination is governed by a localized genomic region, but are less tractable in polygenic systems involving numerous loci of moderate effect.

#### Comparison of sex-specific sequencing coverage in quagga mussels

Coverage analyses were conducted in quagga mussel males and females to investigate potential differences in sequencing depth in the candidate sex-linked region. This approach was chosen based on the expectation that coverage would be lower in the sex that differs from the reference genome’s sex (here: male), due to sequence divergence in sex-linked regions. Such divergence can result in reduced mapping efficiency, as sex-specific sequences may not align well to the reference genome. Coverage differences may as well uncover potential signatures of sex-specific duplications or deletions within particular genomic regions. For all coverage analyses, we applied a less stringent filter of the initial raw VCF files to better capture signals of coverage difference between sexes. The following VCFtools parameters were used: --maf 0.01 MISS=0.2, QUAL=30, MIN_DEPTH=6x. Specifically, we did not filter for maximum coverage, we allowed more missing data, indels and multi-allelic SNPs were kept and we applied a less stringent filter for minor allele frequency. We calculated the average coverage per genomic position and per sex using custom bash scripts. Coverage plots and analyses were conducted on the VCF file with the less stringent filter, while all other analyses (*F*_ST_, ν, GWAS, SDpop) were conducted on the VCF file with the more stringent filters.

#### K-mer analysis in quagga mussels

We further performed a k-mer analysis on the quagga mussel dataset to independently verify the presence of a localised XX-XY sex determination system. We identified sex-specific k-mers in the following way: We first excluded one male which had very low sequencing coverage (quagga60_15; SRX21845084). To have the same number of individuals analysed per sex, we excluded one female (quagga1_5; SRX21845082) which had a very high sequencing coverage. For each of the 19 males and 19 females, we counted the number of 31 bp k-mers in their genome from the trimmed read data using Jellyfish v2.2.7. Then k-mers were merged per sex using jellyfish merge and the male and female k-mers were filtered with two different thresholds: the first threshold was a minimum count of 20 per dataset and the second threshold was a minimum count of 40 per dataset. Filtering was necessary to remove noise and reduce the size of the dataset. The different k-mer sets with different filtering values were then compared to obtain: male-specific kmers, female-specific kmers and common k-mers for both filtering values. Then, the four k-mer datasets (male-specific_min20; female-specific_min20; male-specific_min40; female-specific_min40) were mapped back to the quagga mussel reference genome using the mem function of bwa-mem2 v2.2.1 and default parameters. Mapping statistics were retrieved using the flagstat function of samtools v1.20. Since the mapping percentage was higher for the dataset with a minimum count of 40, we further analysed the mapped data from the minimum count of 40. For the male-specific and female-specific datasets, coverage for each position was extracted using the depth function of samtools v1.20 and plotted using R v4.3.0 in Rstudio 2024.12.0+467.

#### Sequence-based characterization of the FoxL2-derived locus within the quagga SD region

During inspection of gene annotations within the quagga mussel SD region, a gene annotated as FoxL2 was identified. Because FoxL2 is a conserved female-associated gene in metazoans, this annotation warranted further sequence-level examination. To enable direct comparison, FoxL2-like sequences were first identified in the quagga mussel genome by BLASTn searches using the zebra mussel FoxL2 sequence as a query. This search revealed multiple *FoxL2*-derived paralogs in the quagga genome, among which the autosomal copy showing the highest sequence similarity to zebra *FoxL2* was designated as the canonical *FoxL2* reference for subsequent analyses.

Targeted sequence comparisons were then conducted between the SD-region *FoxL2*-derived locus (hereafter *FoxL2-Y*) and the canonical *FoxL2* gene (Results). Coding sequences were aligned at the nucleotide and amino-acid levels using MAFFT v7 and inspected for structural variation, including insertions and deletions, premature stop codons, and nonsynonymous substitutions. Conserved protein domains, including the forkhead (FH) domain in the canonical quagga FoxL2 protein, were identified using InterProScan v107.0. Domain annotations were used to assess domain integrity and overall coding potential. The non-homologous intronic region of *FoxL2-Y* was further inspected for repetitive elements using RepeatMasker.

#### Phylogenetic reconstruction of DMRT, Fox, and Sox gene families

Members of the *DMRT*, *Fox*, and *Sox* gene families were identified from the quagga mussel genome using standalone HMMER v3.4. HMM profiles for the three gene families were built from Pfam models for the DM domain (PF00751), forkhead domain (PF00250), and HMG box (PF00505), respectively. Searches were performed against protein-coding gene annotations derived from BRAKER using an E-value threshold of 1e−5. Identified genes were further verified by CD-Search against NCBI’s Conserved Domain Database (CDD) and by BLASTP searches against the NCBI ClusteredNR database. Gene family members from zebra mussel and other representative taxa were identified from Nicolini et al. (*23*) and retrieved from the corresponding databases. Inferred amino acid sequences were aligned with Clustal Omega and automatically trimmed using trimAl. Best-fit models of protein evolution for each alignment were identified with IQ-TREE v3.0.1. Phylogenetic reconstruction was then performed with RAxML-NG using the best-fit model for each gene family. Maximum-likelihood analyses were used to infer the optimal phylogeny, and node support was assessed with 500 bootstrap replicates.

### Transcriptomic screening of FoxL2-Y and FoxL2 using diagnostic k-mers

Based on sequence differences identified between *FoxL2-Y* and *FoxL2*, we designed diagnostic k-mers to distinguish transcripts originating from (i) *FoxL2-Y* in quagga mussels (FoxL2-Y-Q), (ii) canonical *FoxL2* in quagga mussels (FoxL2-Q), and (iii) *FoxL2* in zebra mussels (FoxL2-Z), the latter serving as an internal control for k-mer specificity across mixed-species RNA-seq libraries. For each target sequence, four diagnostic 24-bp k-mers were designed (three sets of four k-mers in total). Within each set, k-mers 1–3 spanned an 18-bp deletion specific to *FoxL2-Y* together with multiple linked single-nucleotide variants in exon 1, and k-mer 4 spanned a *FoxL2-Y*–specific 2-bp deletion together with several linked SNVs in exon 2, enabling discrimination among highly similar *FoxL2*-derived loci. For all k-mers, each target sequence differed from the alternative FoxL2-derived sequences by at least two diagnostic nucleotide differences (ranging from 2 to 20 differences across the 24-bp k-mer), ensuring robust locus-specific detection. The genomic uniqueness of *FoxL2-Y* diagnostic k-mers was verified against the quagga mussel reference genome using BLASTn. Diagnostic k-mers were queried against publicly available embryonic and developmental RNA-seq datasets as well as newly generated adult tissue-specific RNA-seq libraries (see below), using Jellyfish v2.2.7. When *FoxL2-Y*–specific k-mers were detected, the corresponding read pairs were extracted and merged to reconstruct transcript fragments and validate the presence of multiple linked diagnostic variants within individual reads.

#### Embryonic and developmental RNA-seq datasets

Publicly available RNA-seq datasets covering early development in quagga mussels were obtained from a previous study (*24*). These datasets comprised 23 libraries spanning six developmental stages, including unfertilized eggs, gastrulae, trochophores, early veligers, D-shaped veligers, and juvenile stages. The majority of libraries consisted exclusively of quagga mussel material, while zebra mussels were present only in a small subset of mixed-species libraries and in a single whole-juvenile sample. Libraries varied in sequencing depth, ranging from approximately 15 million to over 100 million paired-end reads (Table S6). These datasets were used primarily to assess the temporal expression of *FoxL2-Y* and canonical *FoxL2* in quagga mussels across early development. Detection of zebra mussel *FoxL2* in mixed-species and juvenile libraries served as an internal positive control for k-mer specificity and method performance, rather than as a basis for stage-specific expression analyses in zebra mussels.

#### Adult tissue-specific RNA sequencing in quagga mussels

Adult quagga mussels for RNA sequencing were collected from the forebay of Davis Dam near Bullhead City, Arizona, USA (35.196077, −114.56899) on April 29, 2022. Mussels were maintained in aerated freshwater until dissection. Individuals were sexed under a stereomicroscope, and tissues including digestive gland, foot, gill, mantle, and gonads were dissected using sterile instruments. Tissue samples were preserved in RNAlater (Thermo Fisher Scientific) and stored at −80°C until RNA extraction. Gonadal tissues were split at dissection, with half preserved for histological validation.

RNA was extracted using either the RNeasy Mini Kit (Qiagen) or the PureLink RNA Mini Kit (Thermo Fisher Scientific) following manufacturer protocols. RNA quality was assessed using an Agilent TapeStation, and all samples had RNA integrity numbers (RIN) ≥8.9. A total of 41 samples were selected for sequencing, including multiple somatic tissues and gonads from both sexes. Libraries were prepared using poly(A) selection and sequenced as 150 bp paired-end reads on an Illumina platform, targeting 20–30 million read pairs per library. Final sequencing depth ranged from 17.7 to 31.0 million paired-end reads per sample, with ≥90% of bases achieving quality scores ≥Q30.

## Author contributions

AATW: Conceptualization, Supervision, Project administration, Investigation, Methodology, Formal analysis, Data curation, Visualization, Resources, Writing – original draft, Writing – review & editing. YP: Investigation, Methodology, Formal analysis, Data curation, Resources, Writing – original draft, Writing – review & editing. KMK: Investigation, Methodology, Resources, Writing – original draft, Writing – review & editing. KU: Investigation, Formal analysis, Methodology. MCS: Investigation, Writing – review & editing. MG: Investigation. SGS: Investigation. ZC: Investigation, Formal analysis, Methodology. JS: Investigation, Formal analysis, Methodology, Writing – review & editing. All authors reviewed and approved the final manuscript.

## Acknowledgements

We are grateful to Linda Haltiner and Christoph Walcher for collecting and maintaining the quagga mussels from Lake Constance (Switzerland), and to Silvana Käser for collecting the zebra mussels from Greifensee (Switzerland). We thank Niklaus Zemp for bioinformatic support. Genome resequencing was performed at the Functional Genomics Center Zürich (FGCZ), Switzerland. Data analyses were performed at the Euler High-Performance Computer Centre in Zürich, Switzerland and the University of Alabama HPC cluster. Data produced and analyzed in this paper were generated in collaboration with the Genetic Diversity Centre (GDC), ETH Zürich. MCS was supported in part by a grant from the U.S. Bureau of Reclamation, Department of the Interior, R19AC000021. KK was supported by a grant from the National Science Foundation - Division of Environmental Biology, #1846174. AATW was supported by the Swiss Federal Institute of Aquatic Science and Technology (Eawag). ZC was supported by the Leibniz Association grant PHENOME (P123/2021). This is contribution number #TBD of the Senckenberg Ocean Species Alliance. YP was supported by grants from the U.S. Bureau of Reclamation’s Science and Technology Program, Department of Interior, ST1866 and ST21024.

## Data accessibility

The zebra mussel chromosome-scale reference genome is publicly available: UMN_Dpol_1.0; NCBI accession: GCF_020536995.1 <u>(McCartney et al. 2022)</u>. The Quagga mussel chromosome-scale reference genome is available from NCBI under the accession number JBLFEX000000000. The corresponding HiFi and Illumina data are available on NCBI SRA (SRR30620554-SRR30620577) under the BioProject ID PRJNA1156045. Quagga mussel raw RNAseq data are available on NCBI SRA (SRR30842657-SRR30842703) under BioProject ID PRJNA1156045. Whole-genome resequencing data of the 80 quagga and zebra mussels are available on NCBI SRA (SRX21845081-SRX21845160) under the BioProject ID PRJNA1019642.

