## Supplementary Figures S1-S10 for "Rapid turnover of sex-determining architecture in invasive mussels"

**Table of contents:**

Supplementary figures:

- **Figure S1:** Exploring genomic signatures for a XY sex determining system in zebra mussels. (p.2)
- **Figure S2:** Genome-wide association study (GWAS) for sex in zebra mussels. (p.3)
- **Figure S3:** Exploring genomic signatures for a ZW sex determining system in quagga mussels. (p.4)
- **Figure S4:** Genome-wide association study (GWAS) for sex in quagga mussels. (p.5)
- **Figure S5:** Nucleotide alignment of coding exons from zebra mussel *FoxL2*, quagga mussel FoxL2, and quagga mussel *FoxL2-Y*, with corresponding quagga embryonic RNA-seq reads. (p.6)
- **Figure S6:** Amino acid alignment of zebra mussel FoxL2, quagga mussel FoxL2, and quagga mussel FoxL2-Y, with translations of corresponding quagga embryonic RNA-seq reads. (p.8)
- **Figure S7:** Chart of gene copy numbers for *DMRT*, *Fox*, and *Sox* gene families. (p.9)
- **Figure S8:** Phylogenetic reconstruction of *Fox* gene family members. (p.10)
- **Figure S9:** Phylogenetic reconstruction of *Sox* gene family members. (p.11)
- **Figure S10:** Phylogenetic reconstruction of *DMRT* gene family members. (p.12)

Supplementary tables (in a separated excel file):

- **Table S1**: individual metadata and statistics of the whole-genome resequencing data of 40 quagga and 40 zebra mussels.
- **Table S2:** list of gene models in the candidate sex-linked regions of zebra mussels.
- **Table S3:** male and female-specific kmer statistics. Dataset of 19 males and 19 females of quagga mussels. kmer length: 31 bp.
- **Table S4:** chromosome-wide average coverage of male-specific and female-specific kmers in quagga mussels. Kmer filtering: minimum 40 counts.
- **Table S5:** list of gene models and their putative function in the candidate XY determining region of quagga mussels.
- **Table S6:** Manual curation of the Y-linked *FoxL2*-derived locus (*FoxL2-Y*) in the quagga mussel sex-determining region.
- **Table S7:** Counts of diagnostic FoxL2-Y-Q, FoxL2-Q, and FoxL2-Z k-mers across embryonic and developmental RNA-seq libraries.
- **Table S8:** Counts of diagnostic FoxL2-Y-Q, FoxL2-Q, and FoxL2-Z k-mers across tissue-specific adult RNA-seq libraries.

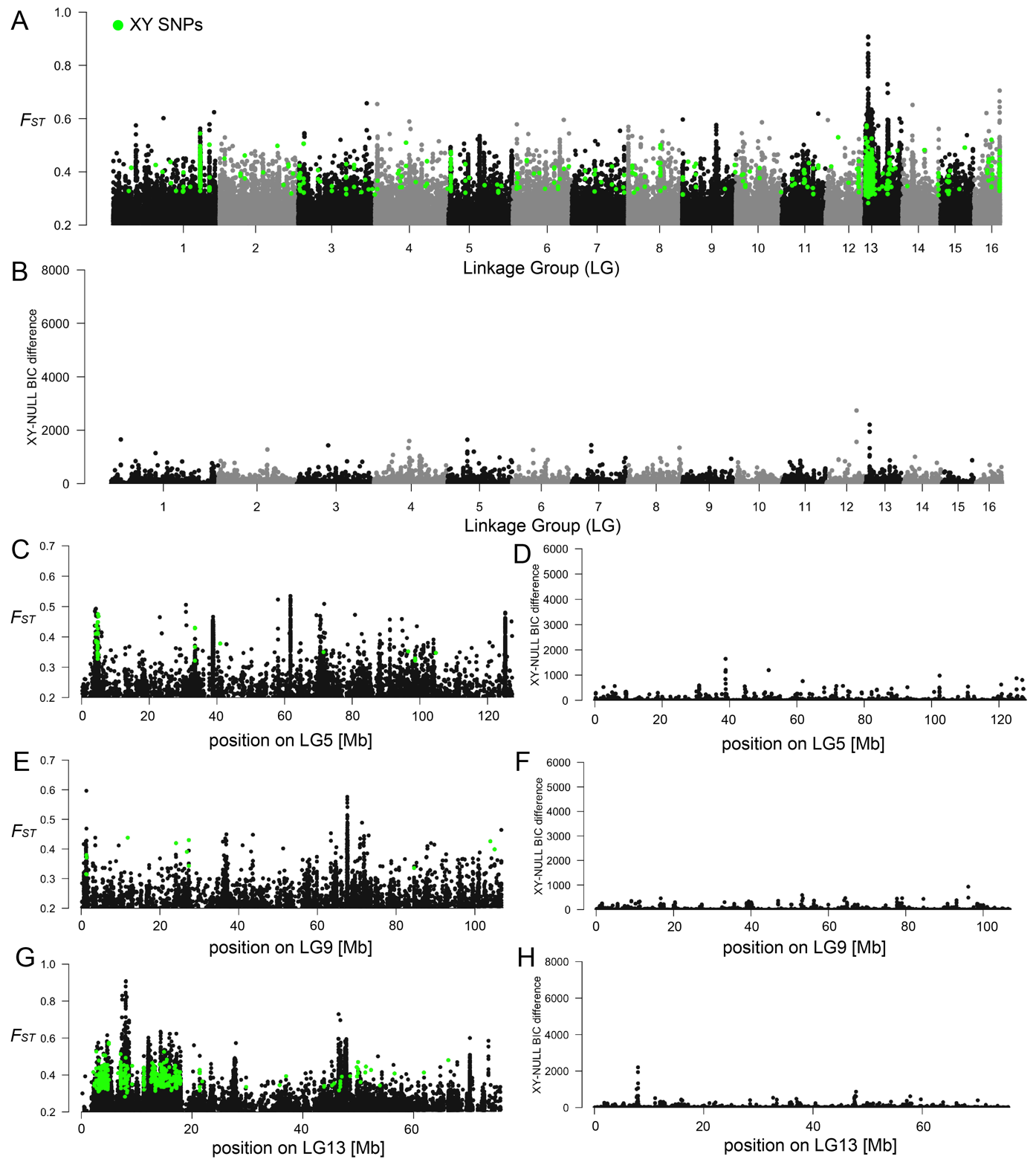

**Figure S1: Exploring genomic signatures for a XY sex determining system in zebra mussels. A)** SNP-based whole-genome Weir & Cockerham *F*_ST_ between 20 male and 20 female zebra mussels. XY SNPs (homozygous in females and heterozygous in males) are highlighted in green. Note that the Y axis starts at 0.2 for better readability in all *F*_ST_ plots. **B)** Bayesian Information Criterion (BIC) difference between XY SDpop model and NULL model calculated over 10 kb sliding windows. An elevated BIC difference (>2000) corresponds to more support for the XY model (XX-XY sex determining system) compared with the NULL model (no genomic differentiation between males and females) for a particular 10 kb window. **C)** Male-female genome-wide *F*_ST_ for linkage group 5 (CM035919.1). **D)** BIC differences between XY and NULL SDpop model for linkage group 5 (CM035919.1). **E)** Male-female genome-wide *F*_ST_ for linkage group 9 (CM035923.1). **F)** BIC differences between XY and NULL SDpop model for linkage group 9 (CM035923.1). **G)** Male-female genome-wide *F*_ST_ for linkage group 13 (CM035927.1). **H)** BIC differences between XY and NULL SDpop model for linkage group 13 (CM035927.1).

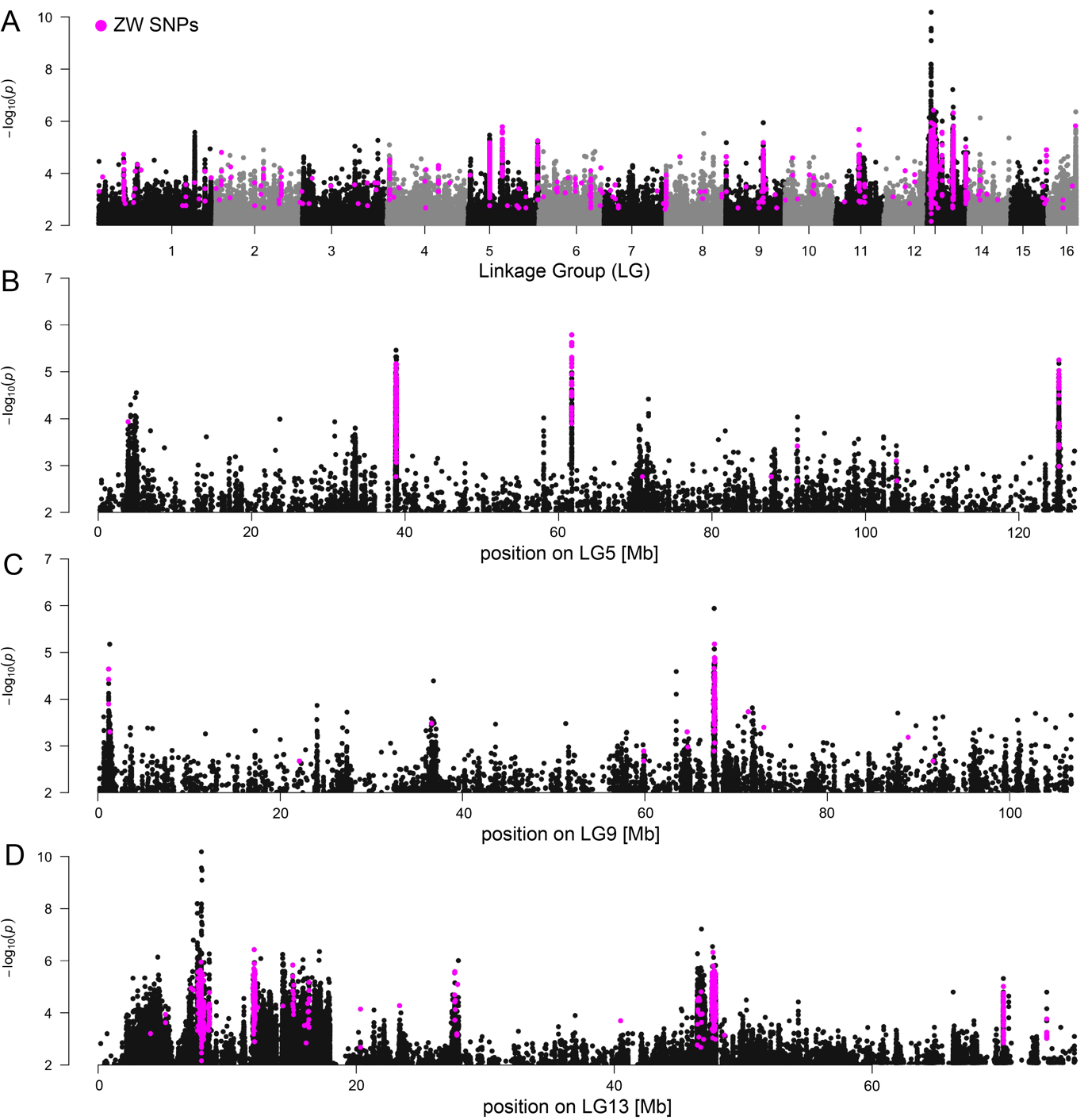

**Figure S2: Genome-wide association study (GWAS) for sex in zebra mussels.** 20 males and 20 females were used. ZW SNPs (homozygous in males and heterozygous in females) are highlighted in pink. Note that the Y axis starts at 2 for better readability in all GWAS plots. **A)** GWAS results across all linkage groups. **B)** GWAS results for linkage group 5. **C)** GWAS results for linkage group 9. **D)** GWAS results for linkage group 13.

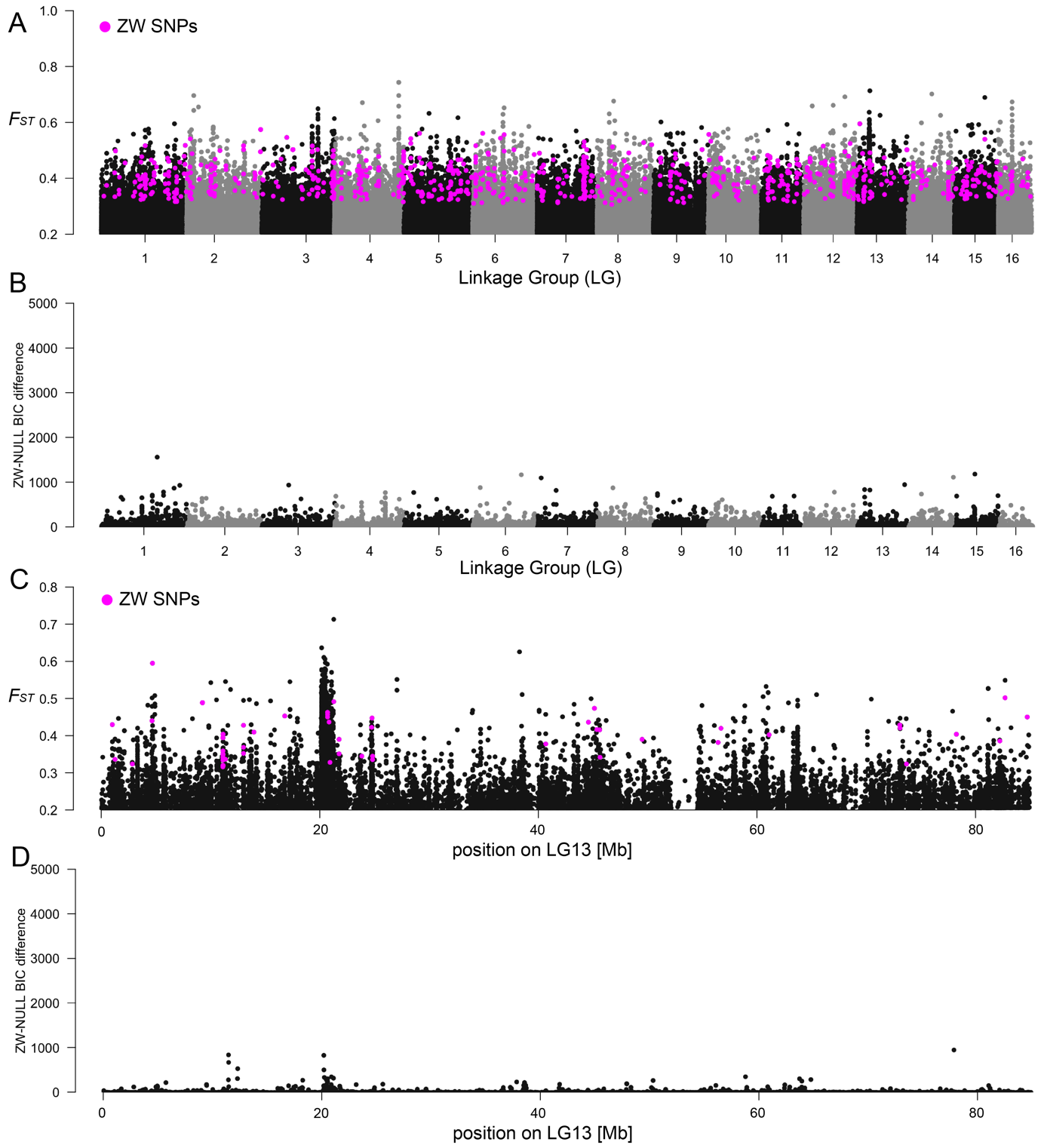

**Figure S3: Exploring genomic signatures for a ZW sex determining system in quagga mussels. A)** SNP-based whole-genome Weir & Cockerham *F*_ST_ between 19 male and 18 female quagga mussels. ZW SNPs (heterozygous in females and homozygous in males) are highlighted in pink. Note that the Y axis starts at 0.2 for better readability in all *F*_ST_ plots. **B)** Bayesian Information Criterion (BIC) difference between ZW SDpop model and NULL model calculated over 10 kb sliding windows. An elevated BIC difference (>2000) corresponds to more support for the ZW model (ZZ-ZW sex determining system) compared with the NULL model (no differentiation between males and females) for a particular 10 kb window. **C)** Male-female genome-wide *F*_ST_ for linkage group 13 (scaffold12). **D)** BIC differences between ZW and NULL SDpop model for linkage group 13 (scaffold12).

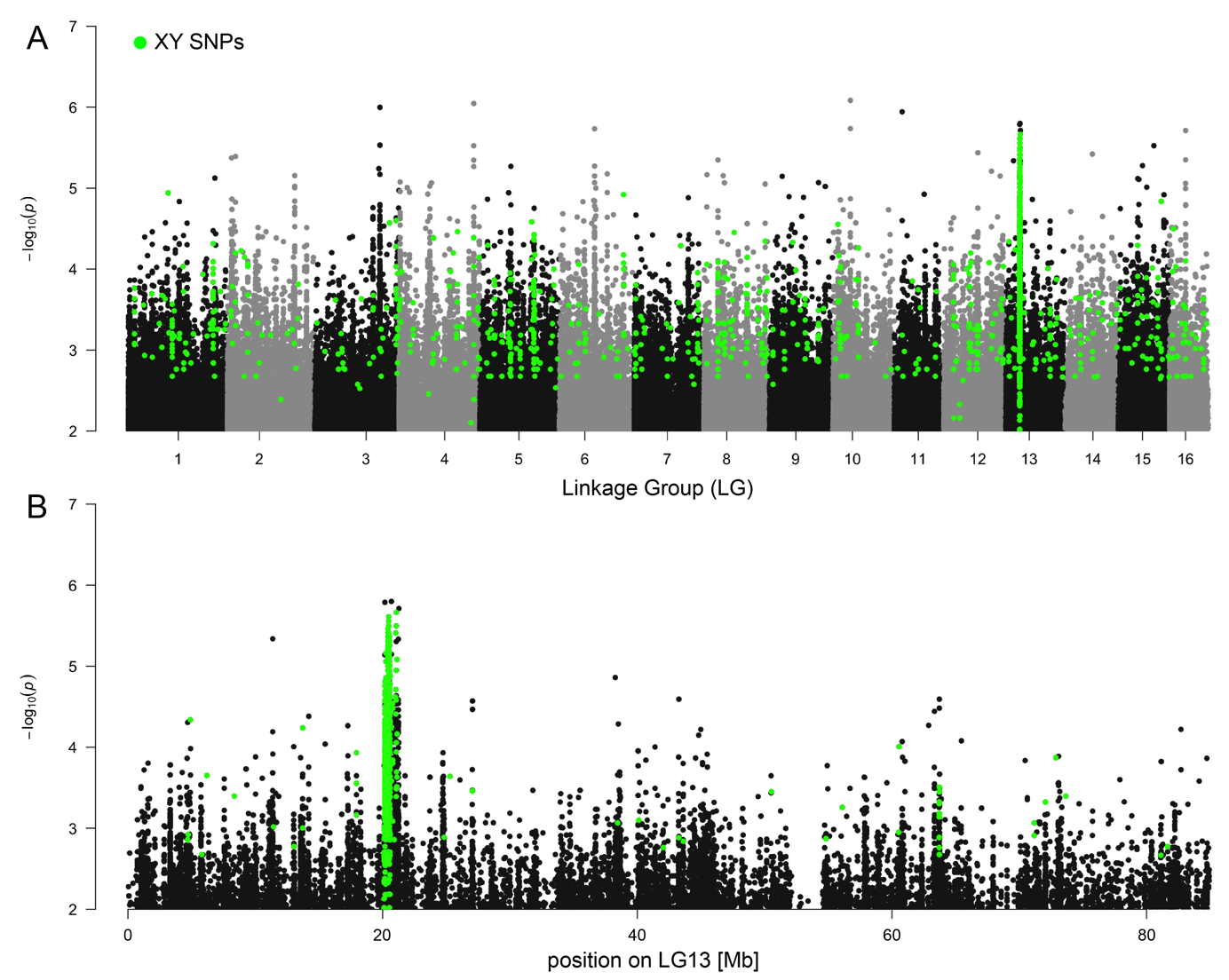

**Figure S4: Genome-wide association study (GWAS) for sex in quagga mussels.** 19 males and 18 females were used. XY SNPs (homozygous in females and heterozygous in males) are highlighted in green. Note that the Y axis starts at 2 for better readability in both GWAS plots. **A)** GWAS results across all linkage groups. **B)** GWAS results for linkage group 13.

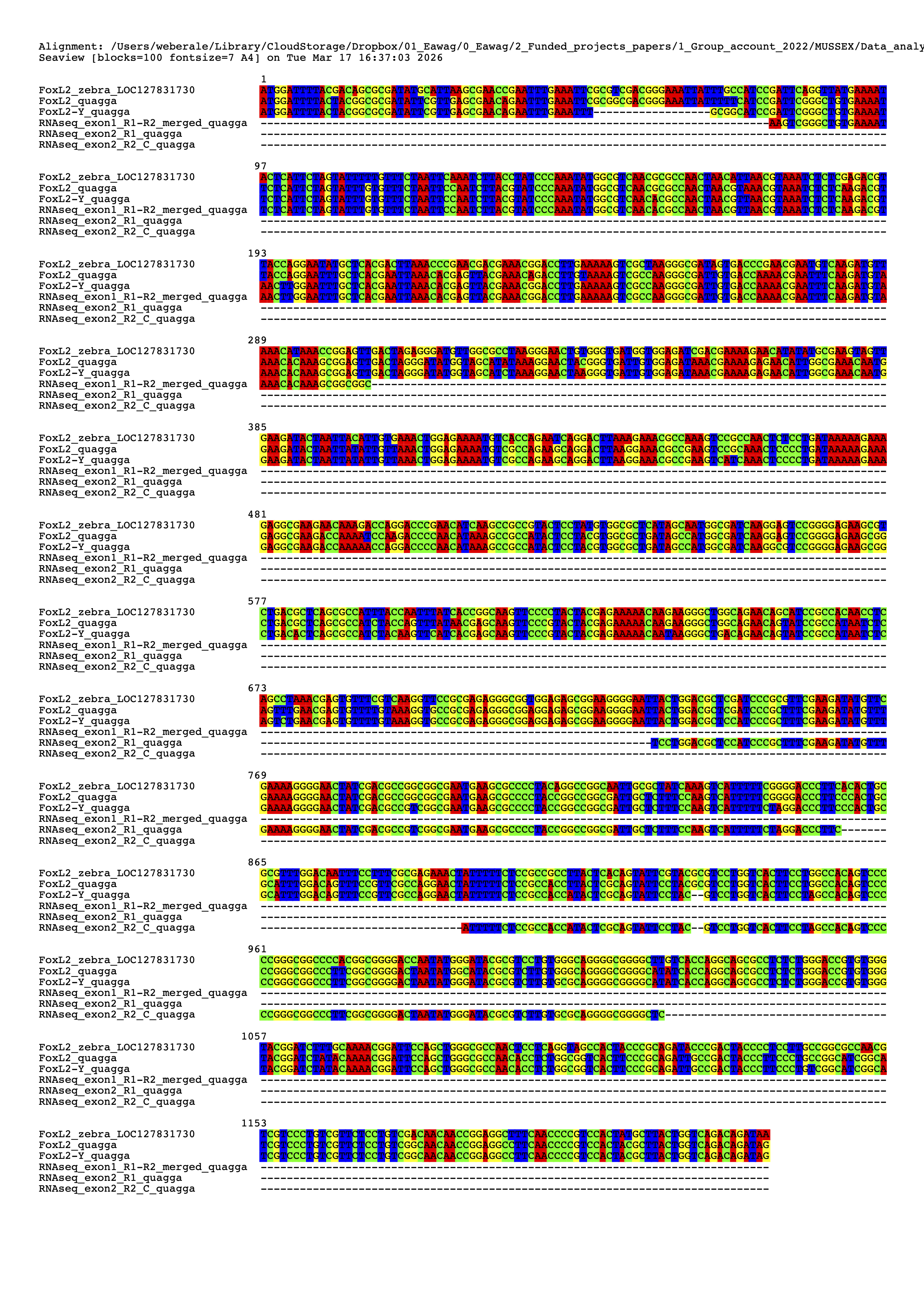

**Figure S5:** **Nucleotide alignment of coding exons from zebra mussel *FoxL2*, quagga mussel *FoxL2*, and quagga mussel *FoxL2-Y*, with corresponding quagga embryonic RNA-seq reads.**

Alignment of exon-only nucleotide sequences from zebra mussel *FoxL2*, quagga mussel *FoxL2*, and the quagga mussel male-specific paralog *FoxL2-Y*. The two lower sequences correspond to the rare embryonic RNA-seq reads recovered from quagga mussel developmental transcriptomes and align specifically to exonic regions of *FoxL2-Y*. Gaps are shown as dashes. This alignment illustrates sequence similarity among the three coding regions while highlighting nucleotide differences distinguishing *FoxL2-Y* from the canonical *FoxL2* copies.

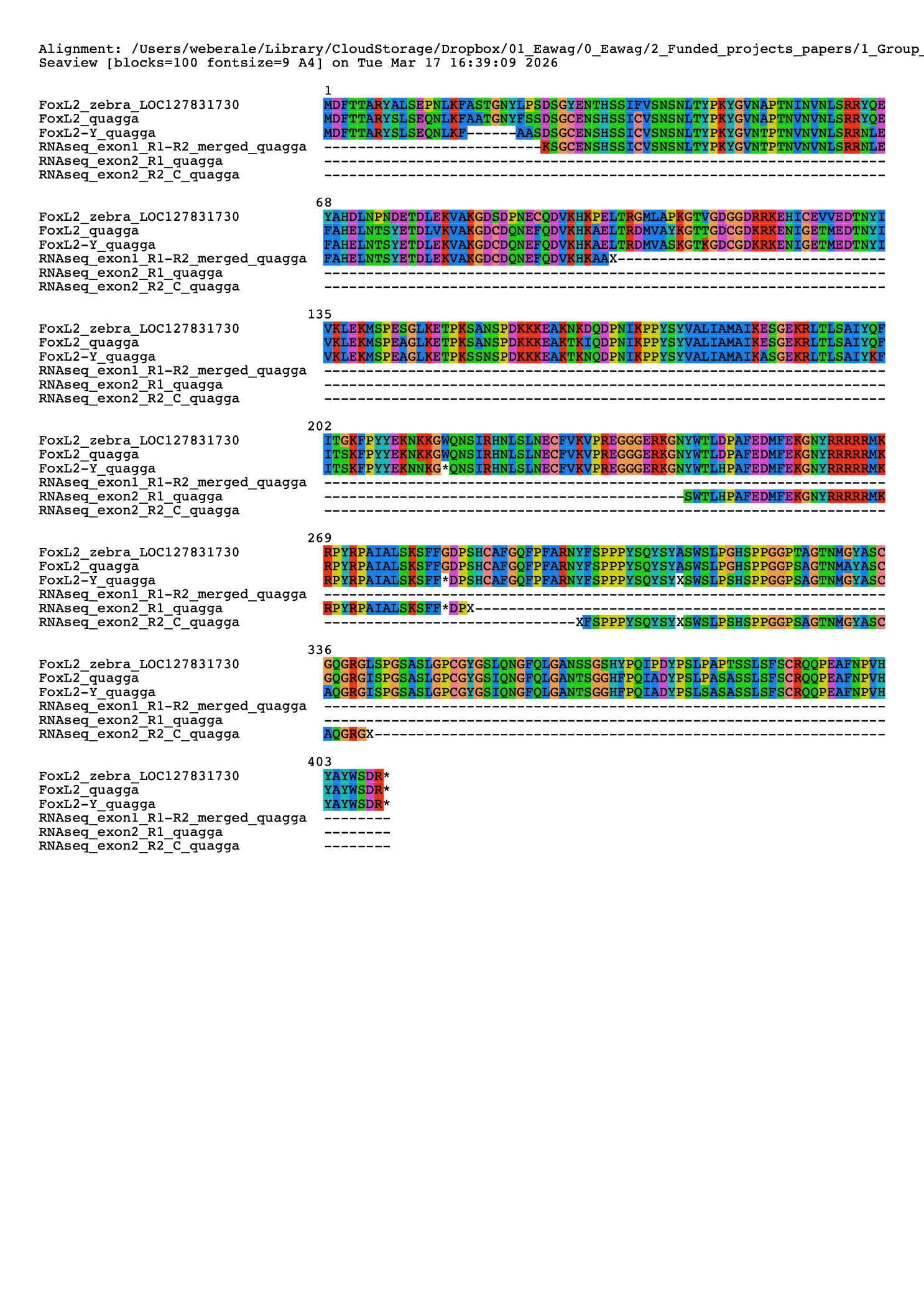

**Figure S6:** **Amino acid alignment of zebra mussel *FoxL2*, quagga mussel *FoxL2*, and quagga mussel *FoxL2-Y*, with translations of corresponding quagga embryonic RNA-seq reads.**

Alignment of predicted amino acid sequences derived from exon-only coding regions of zebra mussel *FoxL2*, quagga mussel *FoxL2*, and quagga mussel *FoxL2-Y*. The two lower sequences represent translations of the rare quagga embryonic RNA-seq reads that map to *FoxL2-Y* exons. Gaps are shown as dashes, and stop codons are indicated by asterisks. The alignment shows that *FoxL2-Y* retains similarity to canonical *FoxL2* across much of the coding sequence but is truncated before the conserved forkhead domain is completed.

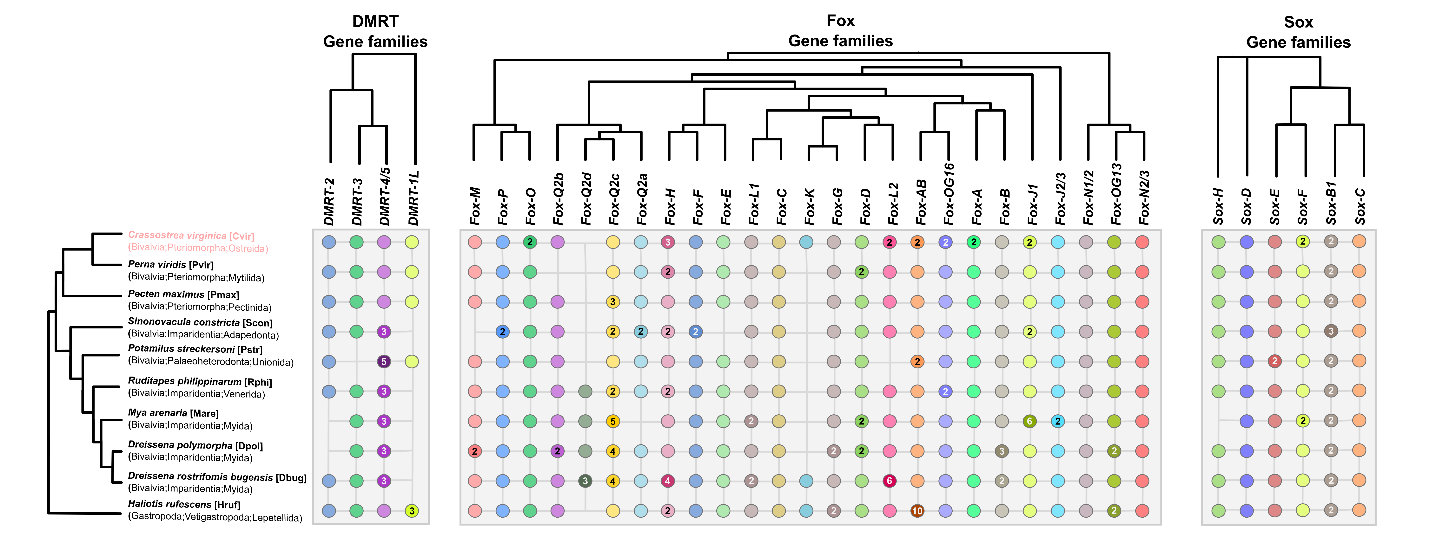

**Figure S7:** **Chart of gene copy numbers for *DMRT*, *Fox*, and *Sox* gene families.** Simplified topologies of gene phylogenies for *DMRT*, *Fox*, and *Sox* are shown along the top of the chart with gene families names for each column. A species phylogeny is shown along the left side of the chart, with species name, along with abbreviations used in the gene phylogenies, and major taxonomic clades for the species on each row. Circles or blank spaces represent presence/absence of a gene family ortholog for a given species. Colors match those used to highlight gene families in the full gene phylogenies. For gene families where multiple paralogs were identified with a species, the number of copies is denoted, and the circle is correspondingly darkened.

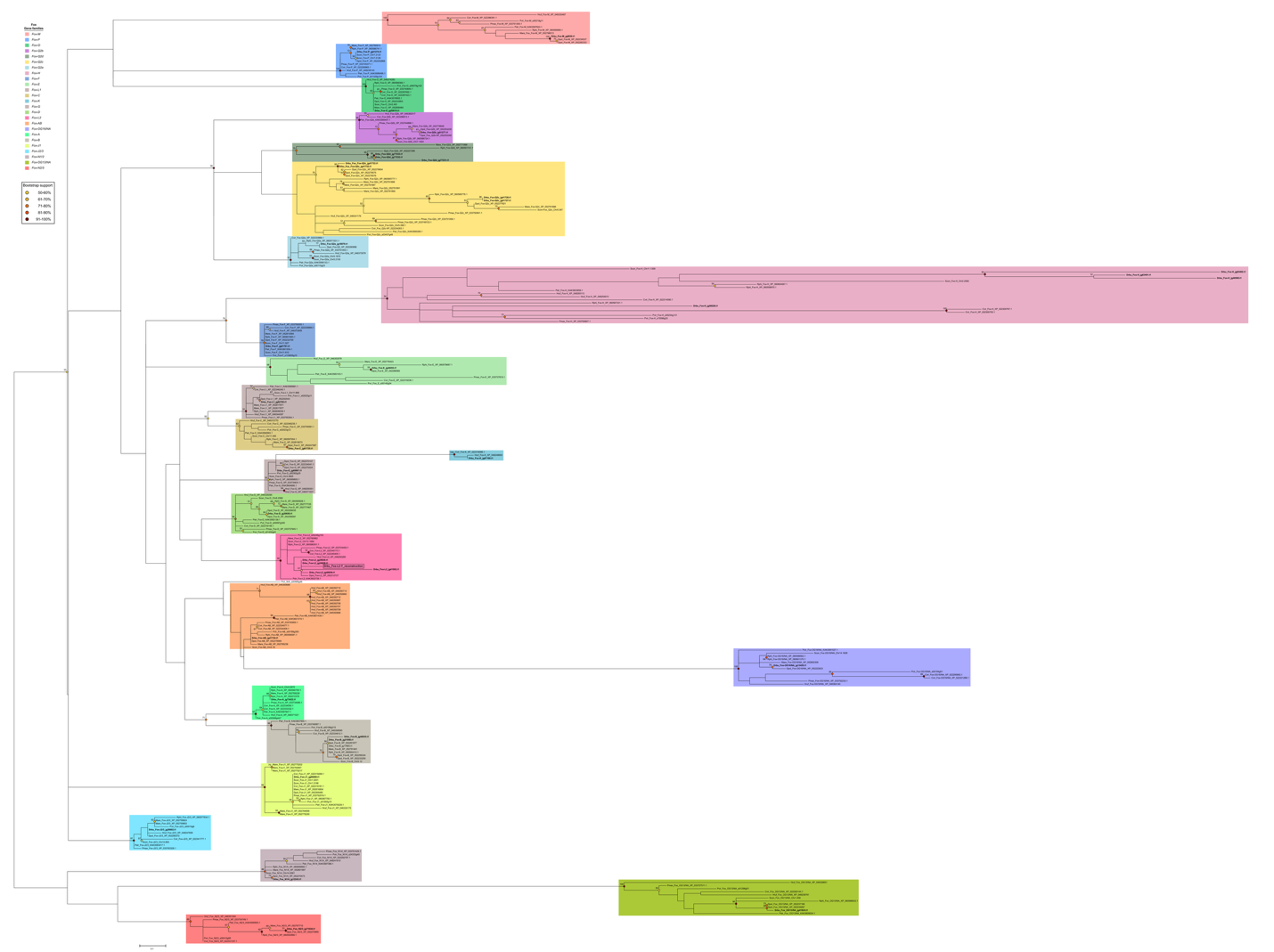

**Figure S8:** **Phylogenetic reconstruction of *Fox* gene family members.** Phylogenetic reconstruction of relationships between *Fox* genes from quagga mussel, zebra mussel, and representative bivalve and gastropod taxa. Gene families are color coded. Nodes with bootstrap support greater than or equal to 50% are color coded, and the bootstrap support is shown adjacent to the node.

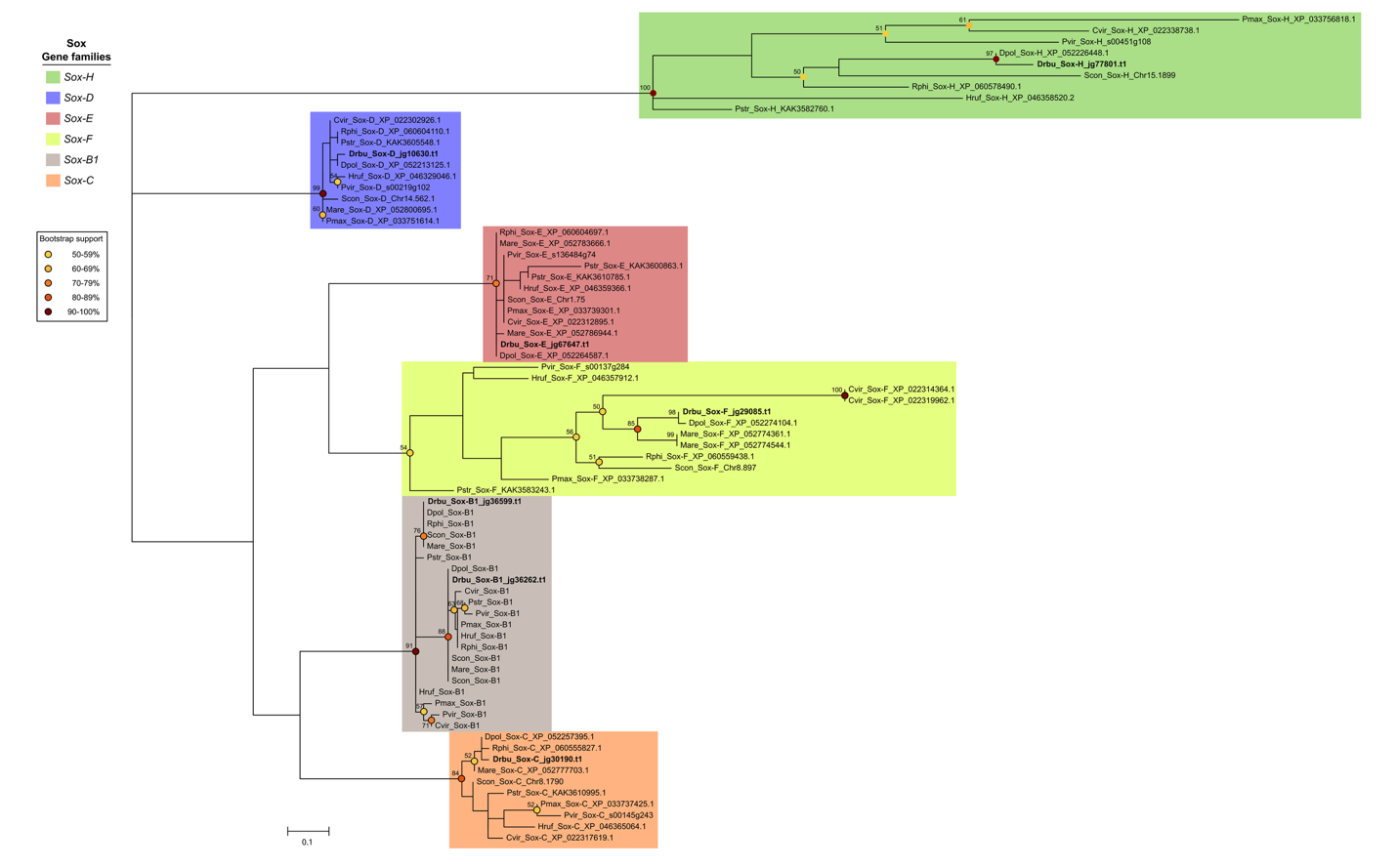

**Figure S9:** **Phylogenetic reconstruction of *Sox* gene family members.** Phylogenetic reconstruction of relationships between *Sox* genes from quagga mussel, zebra mussel, and representative bivalve and gastropod taxa. Gene families are color coded. Nodes with bootstrap support greater than or equal to 50% are color coded, and the bootstrap support is shown adjacent to the node.

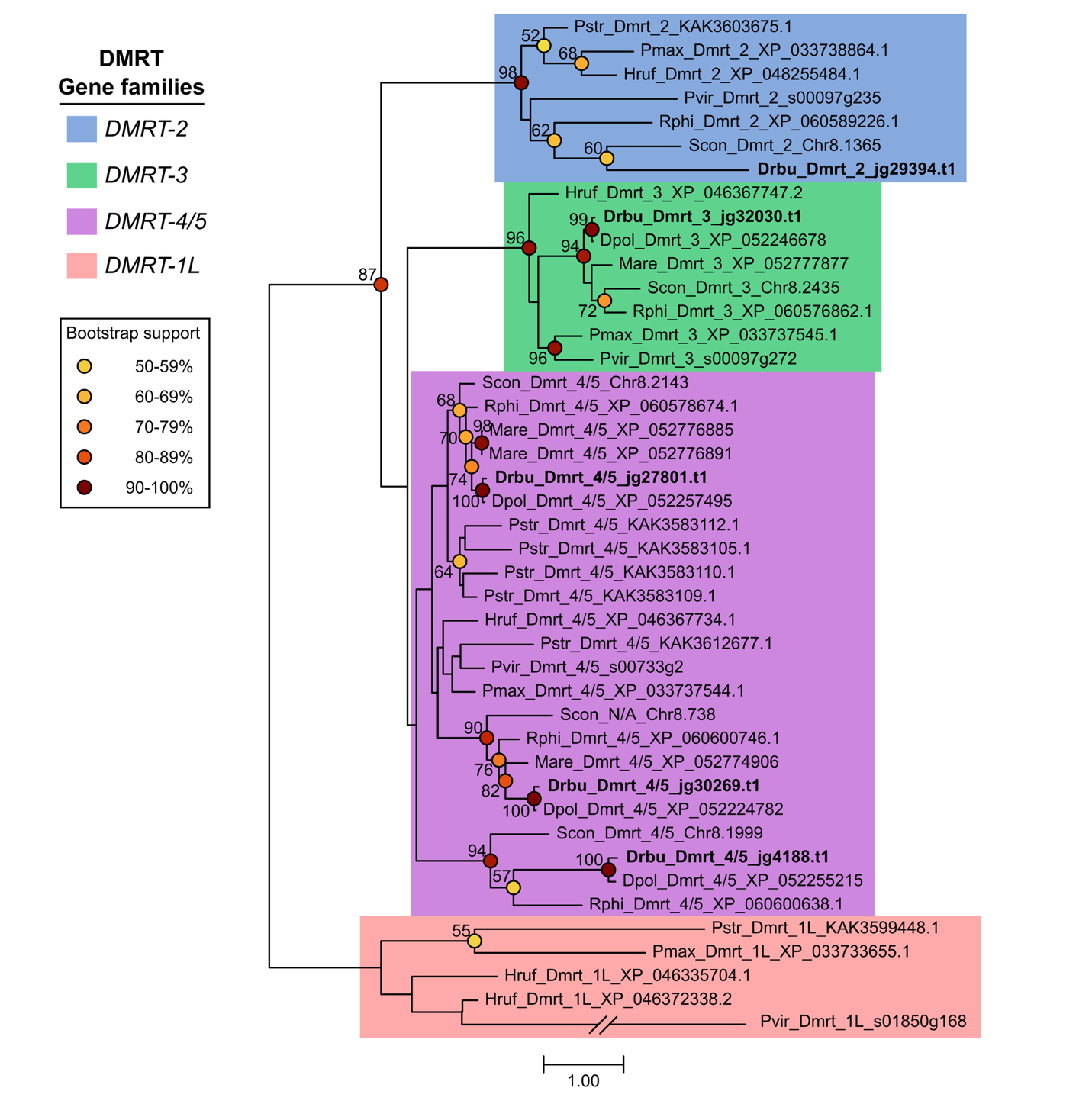

**Figure S10:** **Phylogenetic reconstruction of *DMRT* gene family members.** Phylogenetic reconstruction of relationships between *DMRT* genes from quagga mussel, zebra mussel, and representative bivalve and gastropod taxa. Gene families are color coded. Nodes with bootstrap support greater than or equal to 50% are color coded, and the bootstrap support is shown adjacent to the node.
